# Correlation of *p53* and DNA repair gene mutation patterns in human malignancies indicates different tumour suppression mechanisms of p53

**DOI:** 10.1101/2019.12.19.877068

**Authors:** Xuemin Xue, Lin Dong, Liyan Xue, Yong-jie Lu

## Abstract

P53 suppresses tumorigenesis through multiple cellular functions/mechanisms. Recently, Janic A, et al. reported that DNA repair pathways are critical mediators of p53-dependent tumor suppression. We showed, by mining cBioPortal data of a range of human cancers, that the tendency of ‘mutual exclusivity’ of mutations in *p53* and DNA repair genes only exist in very limited human cancer types. In the majority of human cancers, *p53* mutations are equally distributed between DNA repair gene mutation positive and negative cases and in a number of human cancers, p53 and DNA repair gene mutations have a tendency of co-occurrence. These different correlation patterns of *p53* and DNA repair gene mutations in human malignancies may reflect different critical molecular/cellular pathways activated by p53 in different organs or cell types to suppress tumorigenesis.

---

P53 plays a critical role in suppressing tumor development and is inactivated by gene mutations and/or deletions in half of human cancers^1^. The well-established mechanisms of p53 tumor suppression are induction of cell apoptosis, cell cycle arrest and cell senescence^1^. However, combined loss of cell cycle suspension, apoptosis and senescence did not result in spontaneous tumorigenesis as observed upon loss of p53^2-4^, indicating that there are other critical molecular/cellular mechanisms that p53 activates to suppress tumorigenesis, such as metabolic^2^ and immune response^5,6^ pathways. Exploring the known defective molecular pathways in p53 mutated cancer cells has led to novel forms of tumor therapy strategies^1,7^, thus a complete illustration of all the mechanisms that p53 uses to control tumorigenesis and the specific mechanisms involved in different tumor types should improve cancer therapeutic approaches for *p53* mutated tumors. Recently, Janic et al. reported that DNA repair pathways were critical mediators of p53-dependent tumor suppression^8^.

DNA repair processes are critical for cells to maintain genomic stability. Deficiency in DNA repair processes, frequently caused by DNA repair gene mutations, leads to genomic instability and consequently accumulation of genomic alterations^9^. There are several DNA repair pathways, including mismatch repair, base-excision repair, nucleotide-excision repair, translesion synthesis, homologous recombination, non-homologous end joining, the Fanconi anemia and the O6-methylguanine DNA methyltransferase pathways^9^. Deficiency in mismatch repair pathways leads to microsatellite instability and consequently increased mutation burden and neo-antigen load in tumor cells, which can predict anti-PD-1/PD-L1 immunotherapy response and better than the predicting value of PD-L1 expression level^10^. In the paper published by Janic et al.^8^, the authors demonstrated in mouse models that DNA repair processes are critical mediators of p53-dependent tumor suppression as knockdown of p53 target genes implicated in DNA repair, including Mlh1, Msh2, Rnf144b, Cav1 and Ddit4, accelerated MYC-driven lymphoma development to a similar extent as knockdown of p53, although not all DNA repair genes had equal effect in tumorigenesis. To translate this research finding from mouse models into human cancers, they analyzed leukemia, lymphoma and colorectal cancer data in the cBioPortal and suggested that *p53* and DNA repair gene mutations are mutually exclusive in those human malignancies^8^.

This study provides new insight into p53 tumor suppression mechanisms and would help with the development of new therapeutic approaches. For example, if p53 has a similar role to mismatch repair genes Mlh1 and Msh2, then *p53* mutation, which is common in human cancers, may be used to predict response to anti-PD-1/PD-L1 immunotherapy. We analyzed cBioPortal data^11,12^ of a broad range of human cancers for the general applicability of this association of mutations in *p53* and reported p53-targeted DNA repair genes, as described in the paper by Janic et al^8^. Our reanalysis using the same approach as Janic et al^8^ of the leukemia, lymphoma and colon cancer data produced similar distribution patterns of *p53* mutation in relation to these DNA repair gene mutations^8^. However, apart from colorectal cancer, we did not find an inverse correlation of *p53* and these DNA repair gene mutations (**Supplementary Figure 1**). In hematological malignancies, where the frequencies of mutations in both *p53* and DNA repair genes are very low (each DNA repair gene mutation rate is <1%), the chance of these two types of mutations co-existing in the same patient is expected to be rare. Hence, neither in the original publication (Fig S19)^8^ nor in our analysis (**Supplementary Figure 1**), is mutual exclusivity of mutations in *p53* and these DNA repair genes statistically significant (all p>0.4). Therefore, the data cannot support mutual exclusivity of p53 and DNA repair gene mutations in hematological malignancies.

As amplification is unlikely to cause loss of function of p53 or DNA repair genes, we did further correlation analysis excluding amplification of these genes (see **Supplementary Methods)**. We also performed the correlation analysis between mutations of *p53* and any of these DNA repair genes in combination to increase the statistical power compared to individual DNA repair genes. With this combination, we still did not find a significant inverse correlation between these two types of mutations in hematological malignancies **(Table 1 and Supplementary Figure 2**).

**Table 1.**
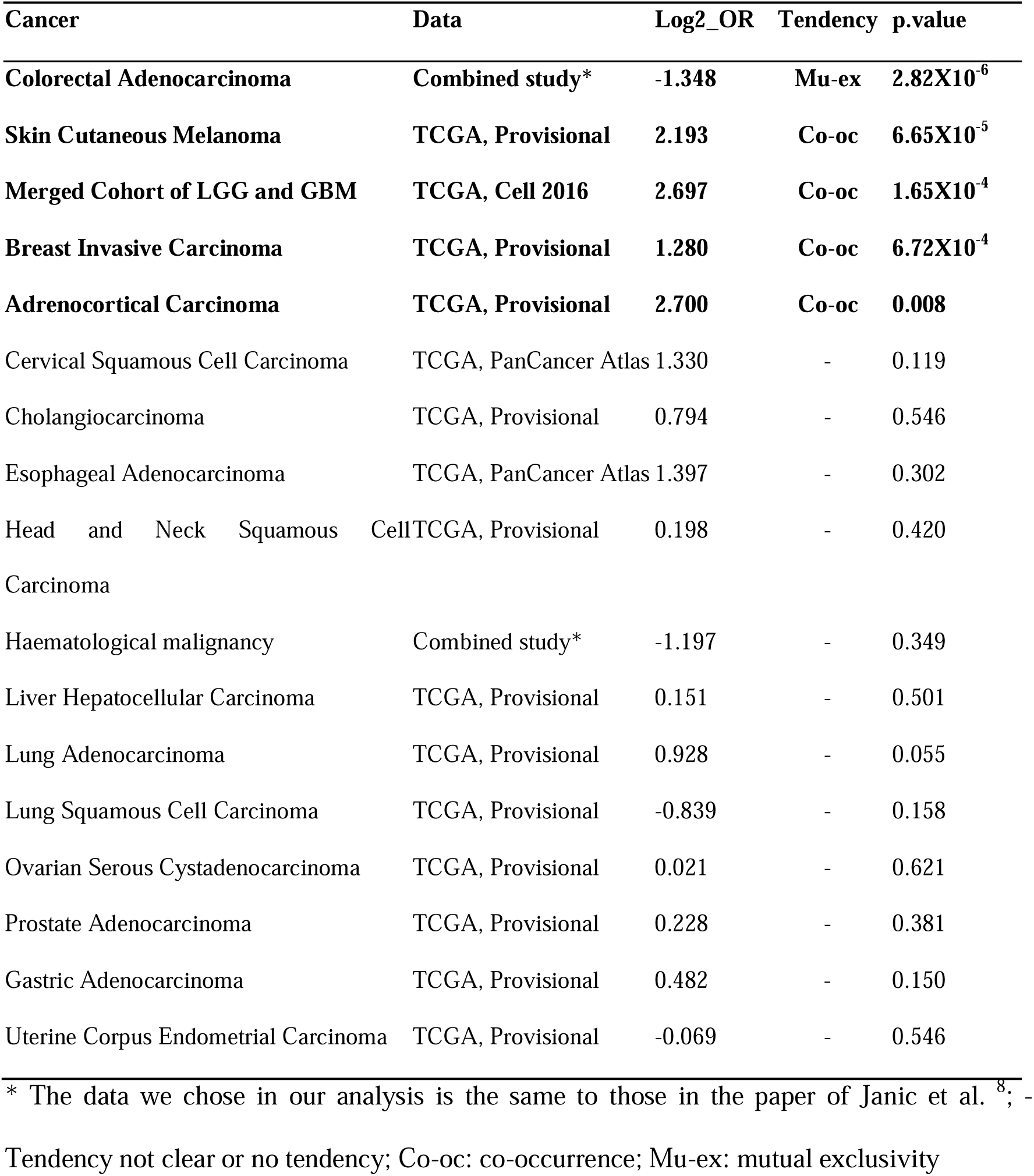
Correlation of mutations in *p53* and any of the 10 p53 regulated DNA repair genes together in human tumors based on genomic sequencing data from cBioPortal^11,12^ datasets

In our further analysis of other human cancers, we found that *p53* mutations are equally distributed between DNA repair gene mutation positive and negative cases in many human cancer types, including prostate, ovarian, liver, head and neck, esophageal, stomach and endometrial cancers and cholangiocarnoma (**Table 1 and Supplementary Figure 2**). Only in lung squamous carcinoma is there a trend (p=0.158) of reverse correlation between *p53* and any of these DNA repair gene mutations with *MLH1* mutation being significantly (p<0.006) reversely correlated with *p53* mutation prior to multiple testing correction. Most importantly, in breast cancer, melanoma, adrenocortical carcinoma and glioma, we found that mutations in *p53* and DNA repair genes are closely associated with each other and these DNA repair gene mutations have a significant (p<0.001, <0.001, 0.008 and <0.001 respectively) tendency of co-occurrence with *p53* mutation (**Table 1, Figure 1**). In cervical squamous carcinoma and lung adenocarcinoma, there is a trend (p=0.119 and 0.055 respectively) of co-occurrence of these two types of mutations, with some DNA repair gene mutations significantly (p<0.05) correlated with *p53* mutation prior to multiple testing adjustment in cervical cancer. In these tumors where *p53* and DNA repair gene mutations have a significant tendency of co-occurrence, DNA repair processes are unlikely to be the mediators of p53-dependent tumor suppression. Therefore, in human malignancies, different correlation patterns of *p53* and DNA repair gene mutations can exist depending on tumor types. The difference in association of *p53* and DNA repair gene mutation patterns may indicate different functions of p53 in action in human malignancies and different therapeutic strategies should be developed accordingly. DNA repair processes may only be the critical mediators of p53-dependent tumor suppression in colorectal cancer and a proportion of *p53* mutated lung squamous carcinoma. In human cancers where *p53* and DNA repair gene mutations co-exist, p53 may play a different role in suppressing tumorigenesis, such as p53-mediated activation of cancer immunity, that p53 function has to be lost to enable cancer cells, with DNA repair gene mutation induced high neo-antigen, to escape the p53-mediated antitumor immunity. This is supported by the observed association between *p53* mutation and lack of tumor infiltrating lymphocytes reported in the literature^6,13^. Further investigations of the p53 functions in these different conditions in association with DNA repair genes are warranted.

**Figure 1.**
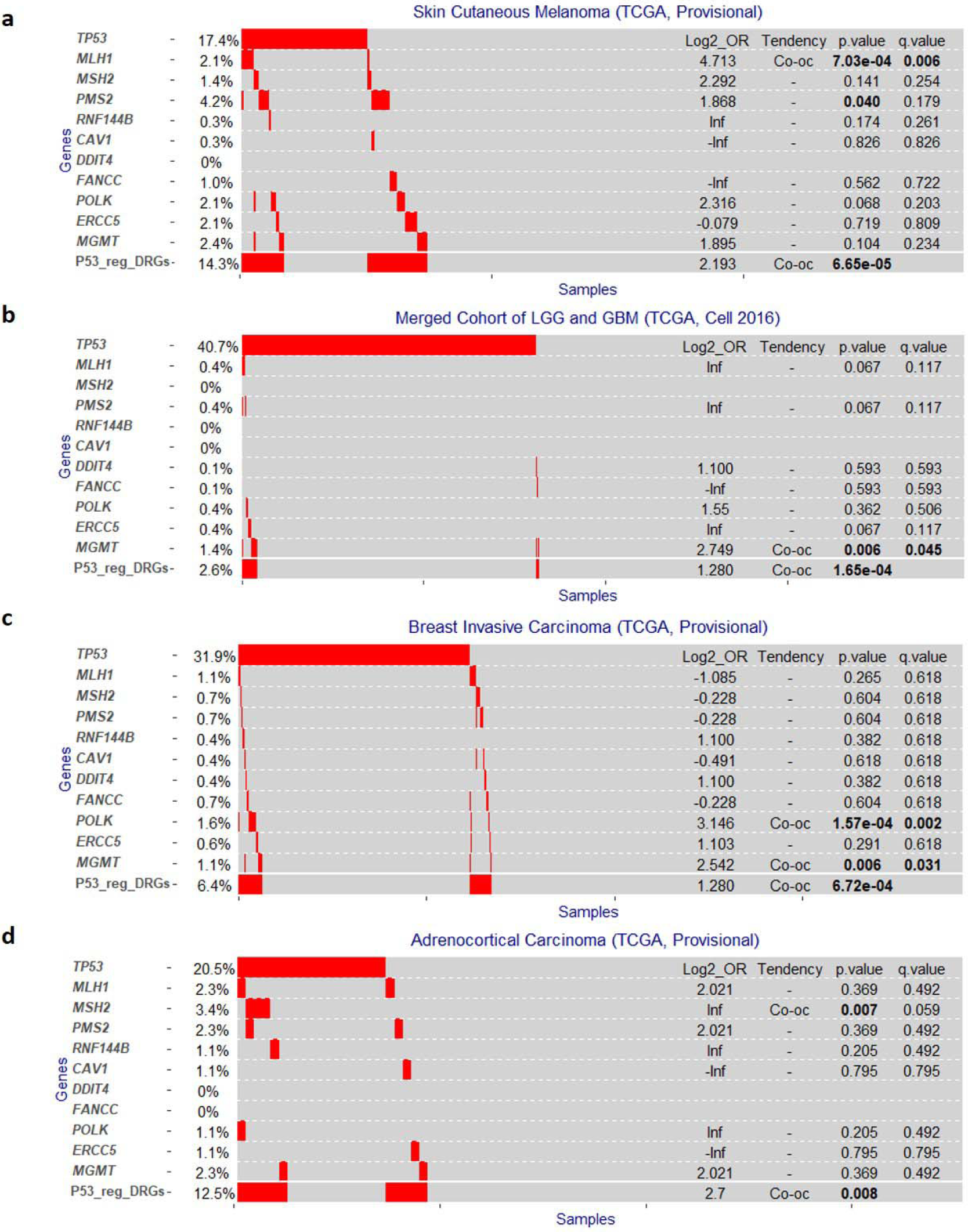
*P53(TP53)* and DNA repair gene mutation distribution in melanoma (a), glioma (b), breast invasive carcinoma (c) and Adrenocortical Carcinoma (d) based on cBioPortal ^11,12^ data. OR: odd ratio; Co-oc: co-occurrence; e-0n: X10^−n^; Inf: Infinity; P53_reg_DRGs: mutation in any of the 10 p53 regulated DNA repair genes in combination.

## Author contributions

YJ.L, and L.X contribute to the concept of the study. X.X and L.D performed the data analysis. YJ.L drafted the manuscript text and X.X, L.D, L.X, and YJ.L wrote the final manuscript.

## Supplementary information

Supplementary methods

Supplementary Figure 1

Supplementary Figure 2

