## Supplemental materials for "Correlation of *p53* and DNA repair gene mutation patterns in human malignancies indicates different tumour suppression mechanisms of p53"

**Supplementary Methods**

1. **Data mining using cBioPortal online tools**

As mentioned in Janic A, et al study^8^, the cBioPortal online data mining were performed at the website <https://www.cbioportal.org> by selecting the dataset(s) of each cancer type and the datasets including number of samples in each study were listed in Supplementary Table 1. Then genes, including *TP53, MLH1, MSH2, PMS2, RNF144B, CAV1, DDIT4, FANCC, POLK, ERCC5* and *MGMT,* were submitted for querying. Heatmap were displayed by clicking on “OncoPrint”. The correlations between genomic alterations of *p53* and these DNA repair genes were generated by clicking on “Mutual Exclusivity”.

2. **Correlation analysis using R statistic software**

The “alterations_across_samples.tsv” file for the dataset(s) of each cancer type was downloaded from cBioPortal website^11,12^. Cases without mutation data (not profiled) were excluded from the correlation analysis (mainly occurred in some datasets for the hematological malignancy and colorectal cancer). The amplifications were also excluded from the genomic alteration data for the analyzed genes. The number of P53_reg_DRGs positive cases were calculated by including samples with mutations in any of the 10 p53 targeted DNA repair genes (*MLH1, MSH2, PMS2, RNF144B, CAV1, DDIT4, FANCC, POLK, ERCC5* and *MGMT)*. Correlation analysis was run using Fisher exact test with one-tail according to cBioPortal^11,12^. P values were adjusted for multiple tests using Benjamini-Hochberg procedure to generate false discovery rate (q value). Tendency of co-occurrence or mutual exclusivity was determined by odd ratio (OR) or Log2_OR^11,12^. Heatmap was plotted using ggplot2 package. All these analysis were run in R 3.4.1 statistic software.

**Supplementary Table 1. Datasets and sample size of each cancer included in the analysis**

| **Cancer types** | **Datasets** | **Sample sizes** |
| --- | --- | --- |
| Colorectal Adenocarcinoma | DFCI, Cell Reports 2016  Genentech, Nature 2012  TCGA, Provisional  MSKCC, Genome Biol 2014  MSKCC, Cancer Cell 2018 | 619  74  640  138  1134 |
| Skin Cutaneous Melanoma | TCGA, Provisional | 287 |
| Merged Cohort of LGG and GBM | TCGA, Cell 2016 | 794 |
| Breast Invasive Carcinoma | TCGA, Provisional | 963 |
| Adrenocortical Carcinoma | TCGA, Provisional | 88 |
| Cervical Squamous Cell Carcinoma | TCGA, PanCancer Atlas | 278 |
| Cholangiocarcinoma | TCGA, Provisional | 35 |
| Esophageal Adenocarcinoma | TCGA, PanCancer Atlas | 182 |
| Head and Neck Squamous Cell Carcinoma | TCGA, Provisional | 504 |
| Haematological malignancy | ALL(St Jude, Nat Genet 2015)  CLL (Broad, Cell 2013)  CLL (IUOPA, Nature 2015)  cTCL (Columbia U, Nat Genet 2015)  DLBCL(Broad, PNAS 2012)  DLBCL (TCGA, Provisional)  MCL (IDIBIPS, PNAS 2013)  MM(Broad, Cancer Cell 2014)  PCNSL (Mayo Clinic, Clin Cancer Res 2015)  AML (TCGA, Provisional)  hALL (St Jude, Nat Genet 2013) | 93  160  506  43  58  48  29  211  19  200  44 |
| Liver Hepatocellular Carcinoma | TCGA, Provisional | 366 |
| Lung Adenocarcinoma | TCGA, Provisional | 230 |
| Lung Squamous Cell Carcinoma | TCGA, Provisional | 178 |
| Ovarian Serous Cystadenocarcinoma | TCGA, Provisional | 311 |
| Prostate Adenocarcinoma | TCGA, Provisional | 492 |
| Gastric Adenocarcinoma | TCGA, Provisional | 393 |
| Uterine Corpus Endometrial Carcinoma | TCGA, Provisional | 242 |

ALL:Acute Lymphoblastic Leukemia; CLL: Chronic Lymphocytic Leukemia; cTCL:Cutaneous T Cell Lymphoma; DLBCL: Diffuse Large B-Cell Lymphoma; MCL: Mantle Cell Lymphoma; MM: Multiple Myeloma; PCNSL: Primary Central Nervous System Lymphoma; AML: Acute Myeloid Leukemia; hALL: Hypodiploid Acute Lymphoid Leukemia.

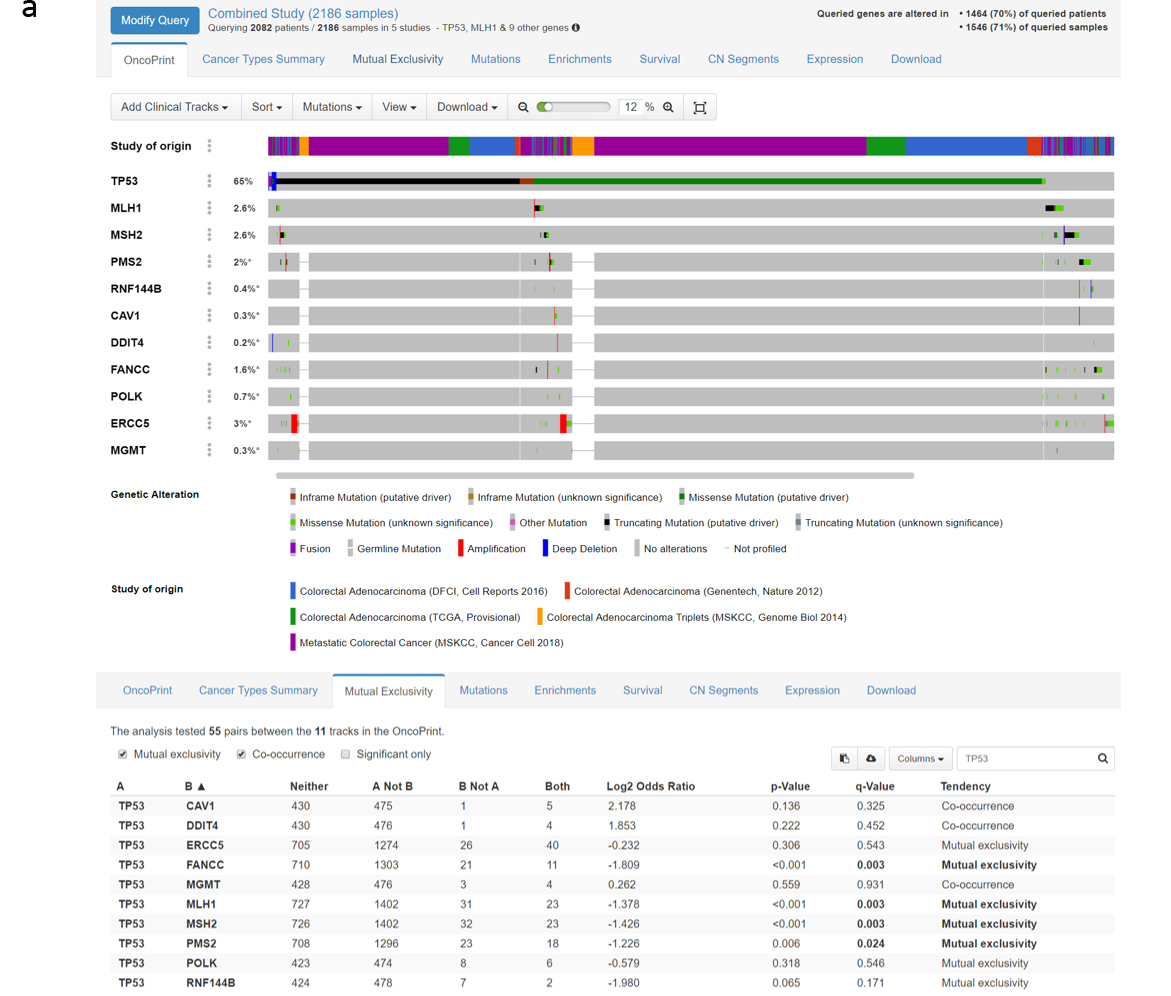

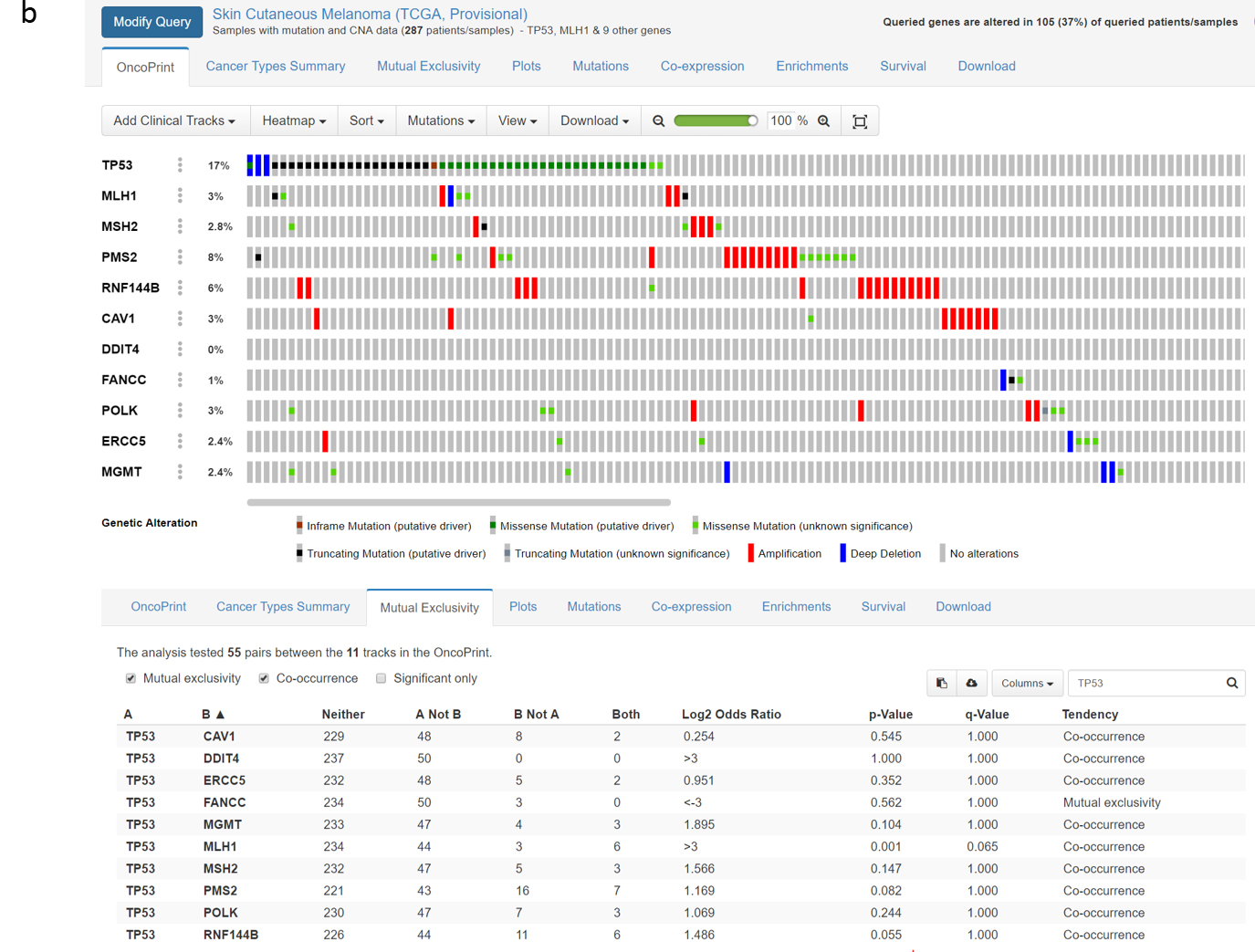

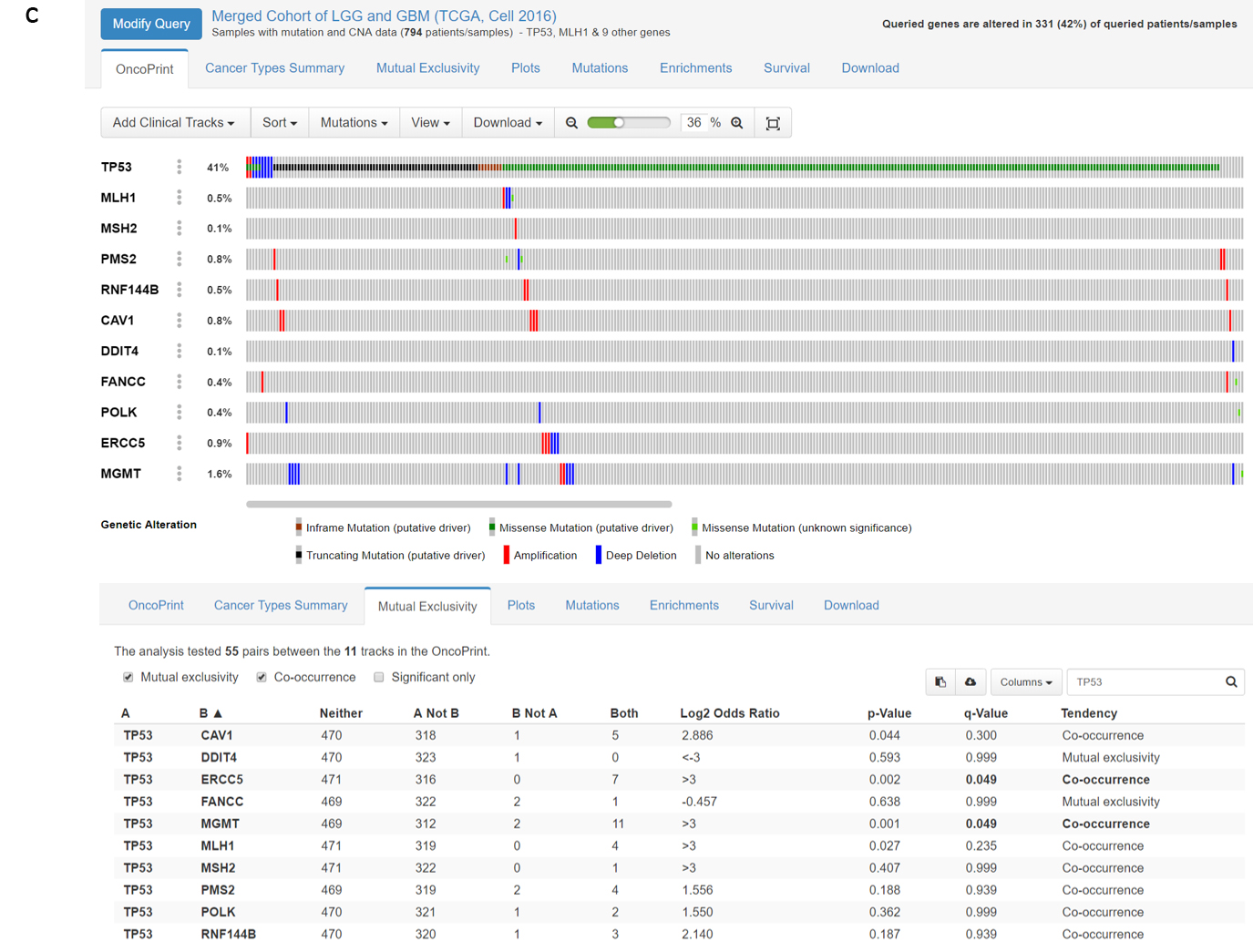

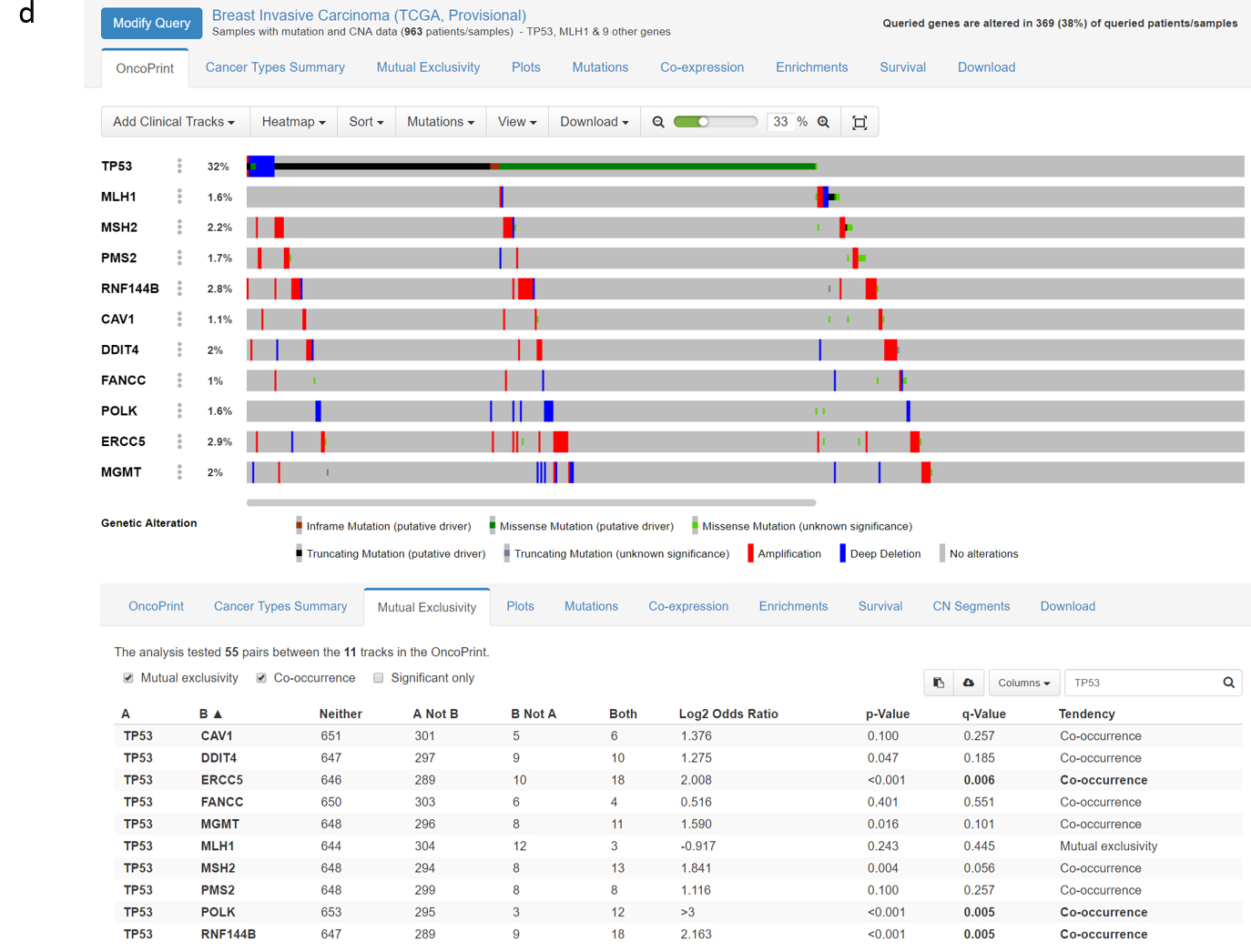

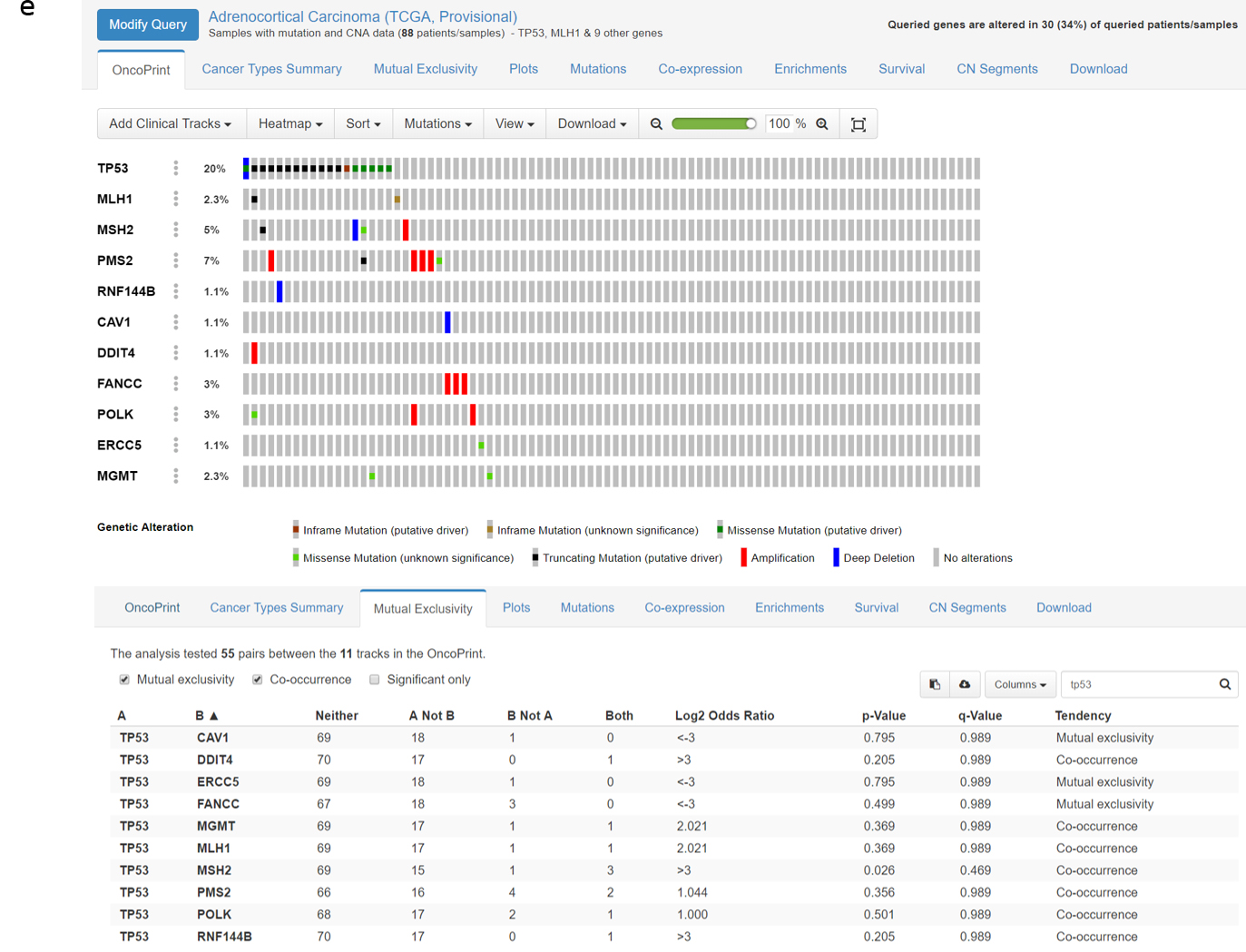

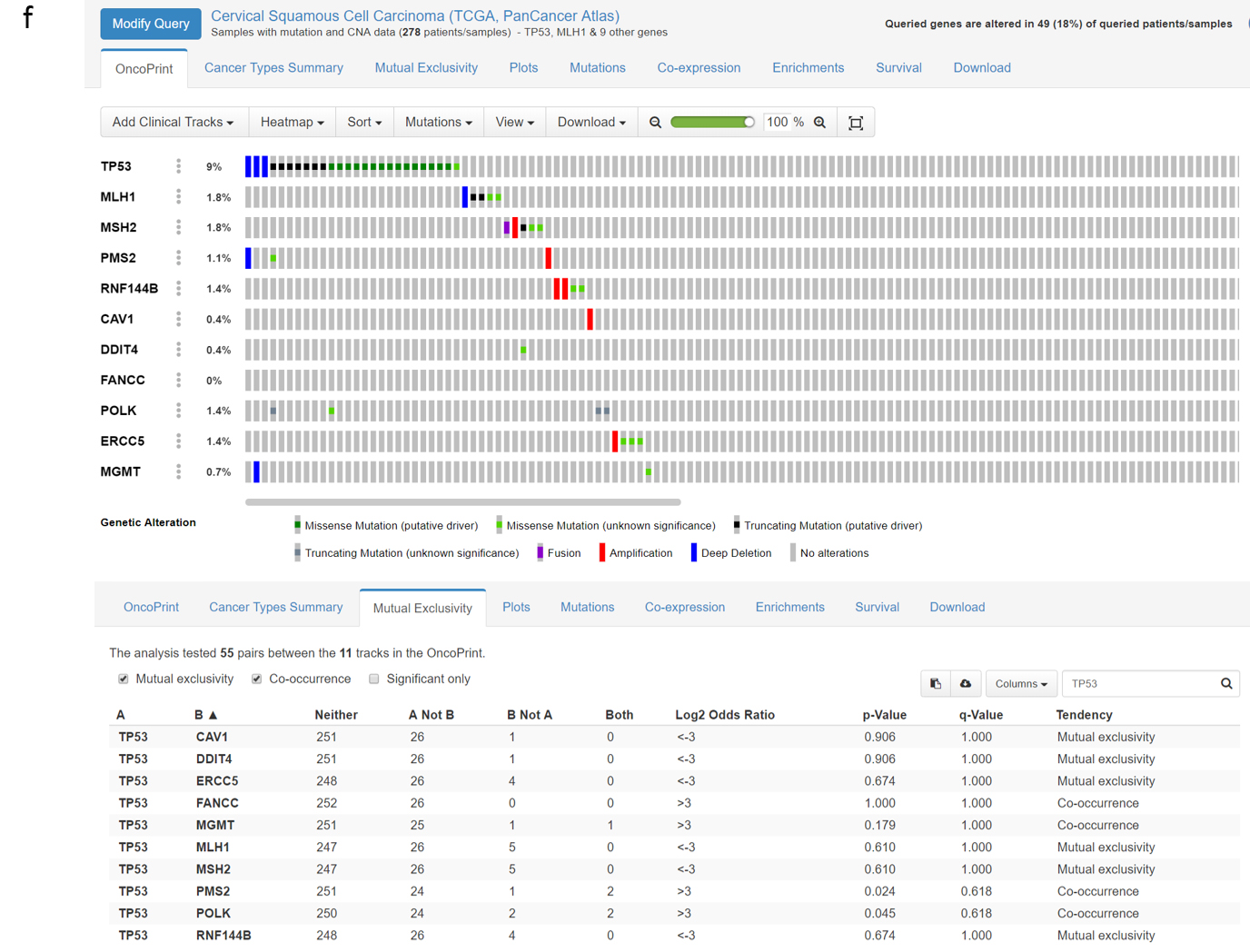

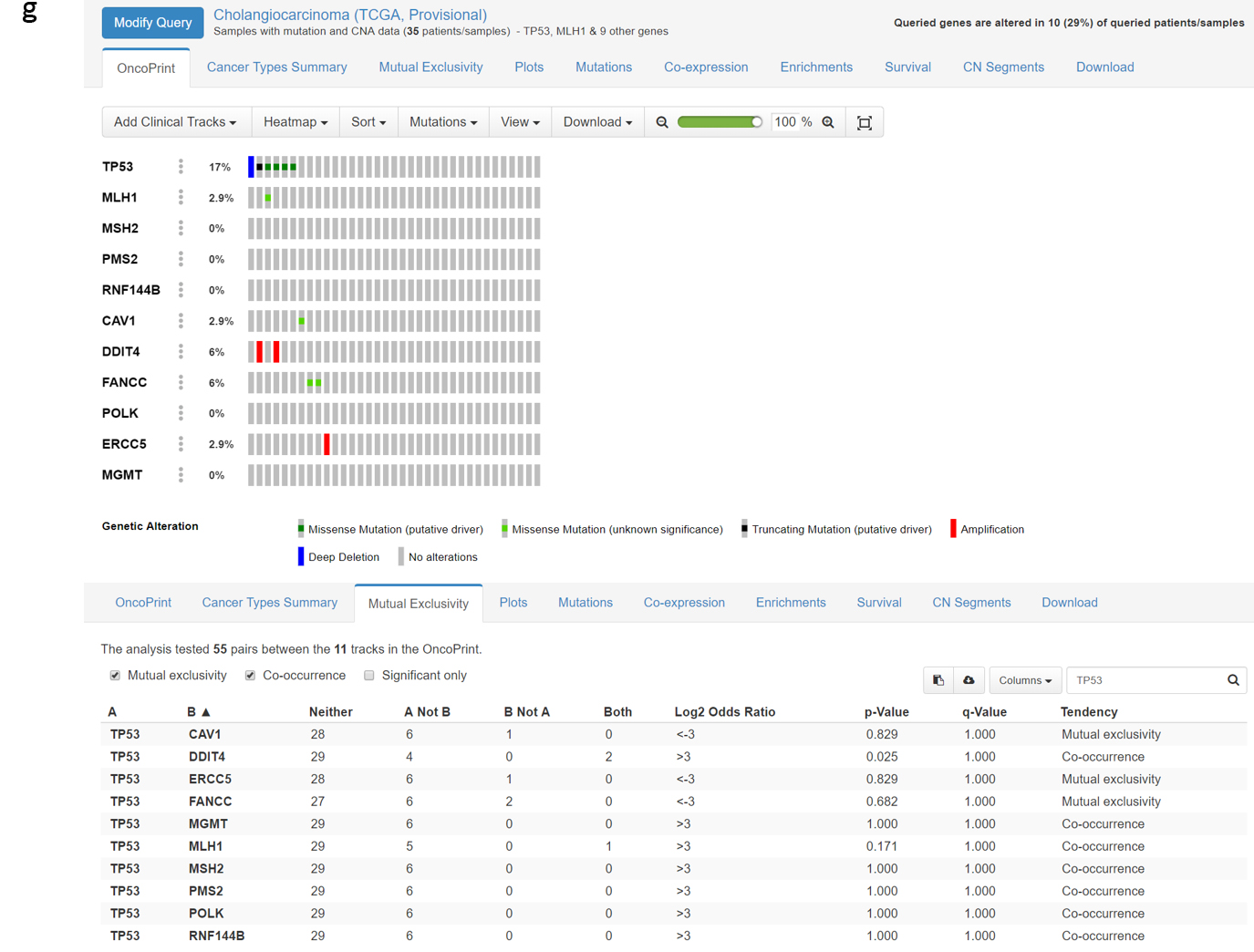

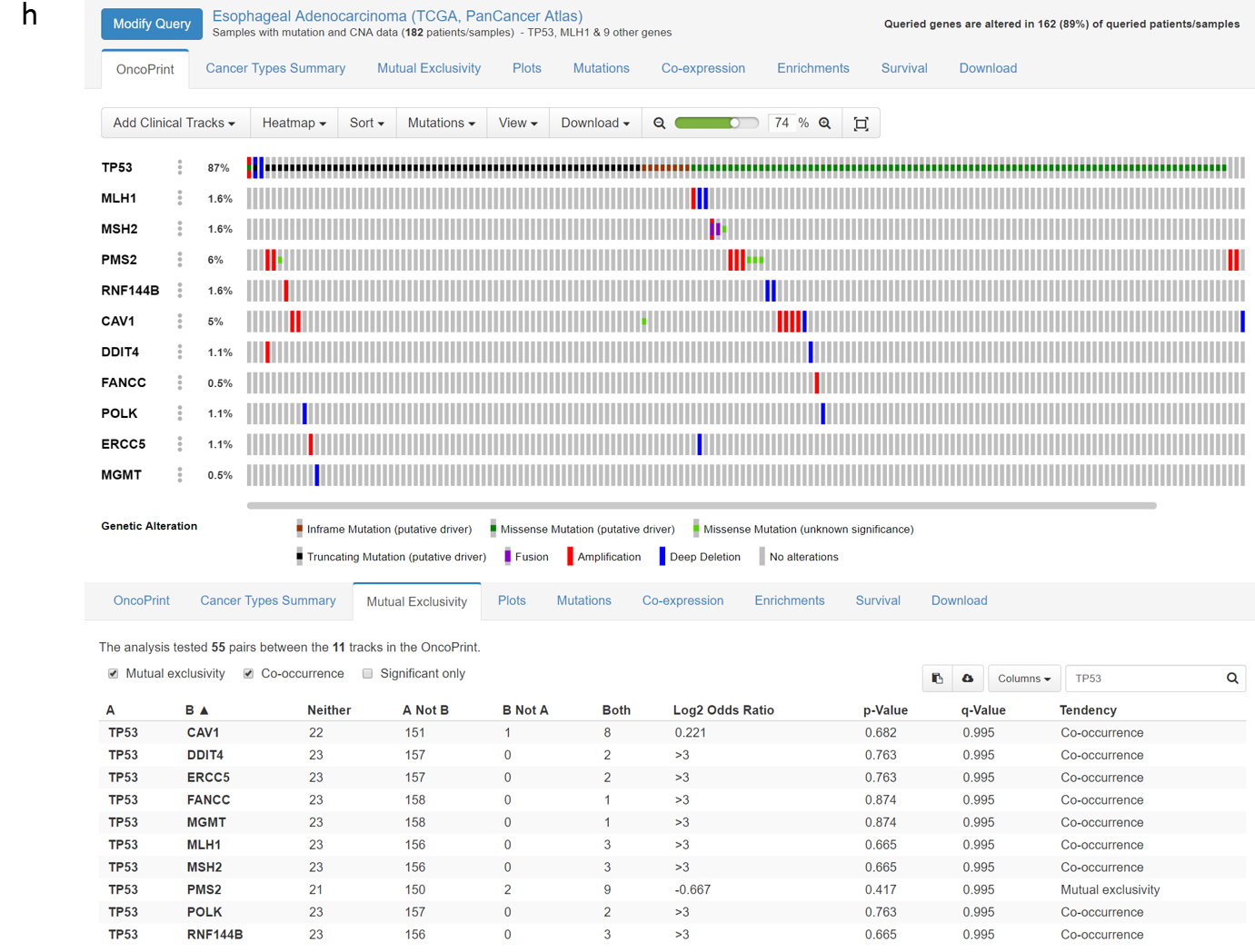

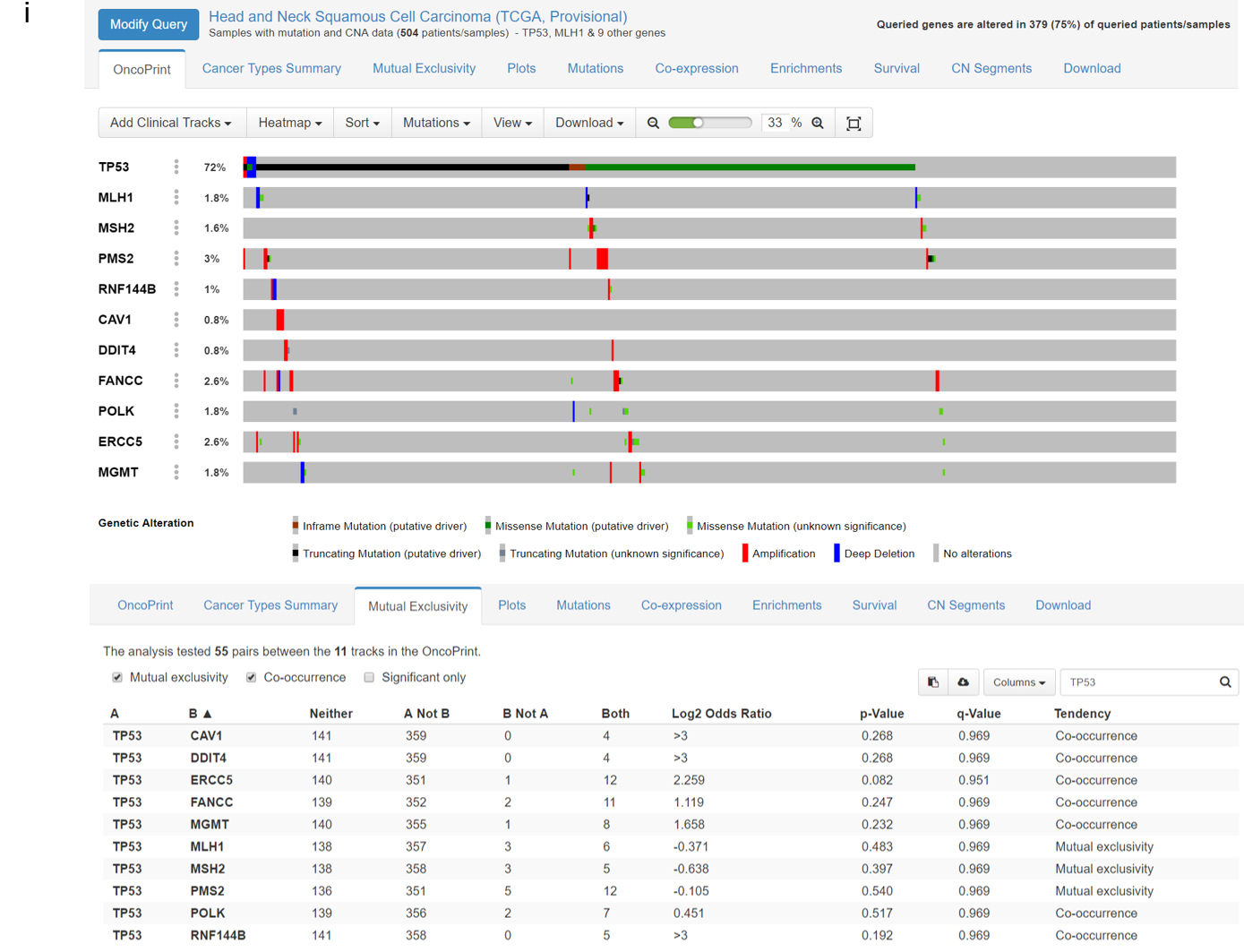

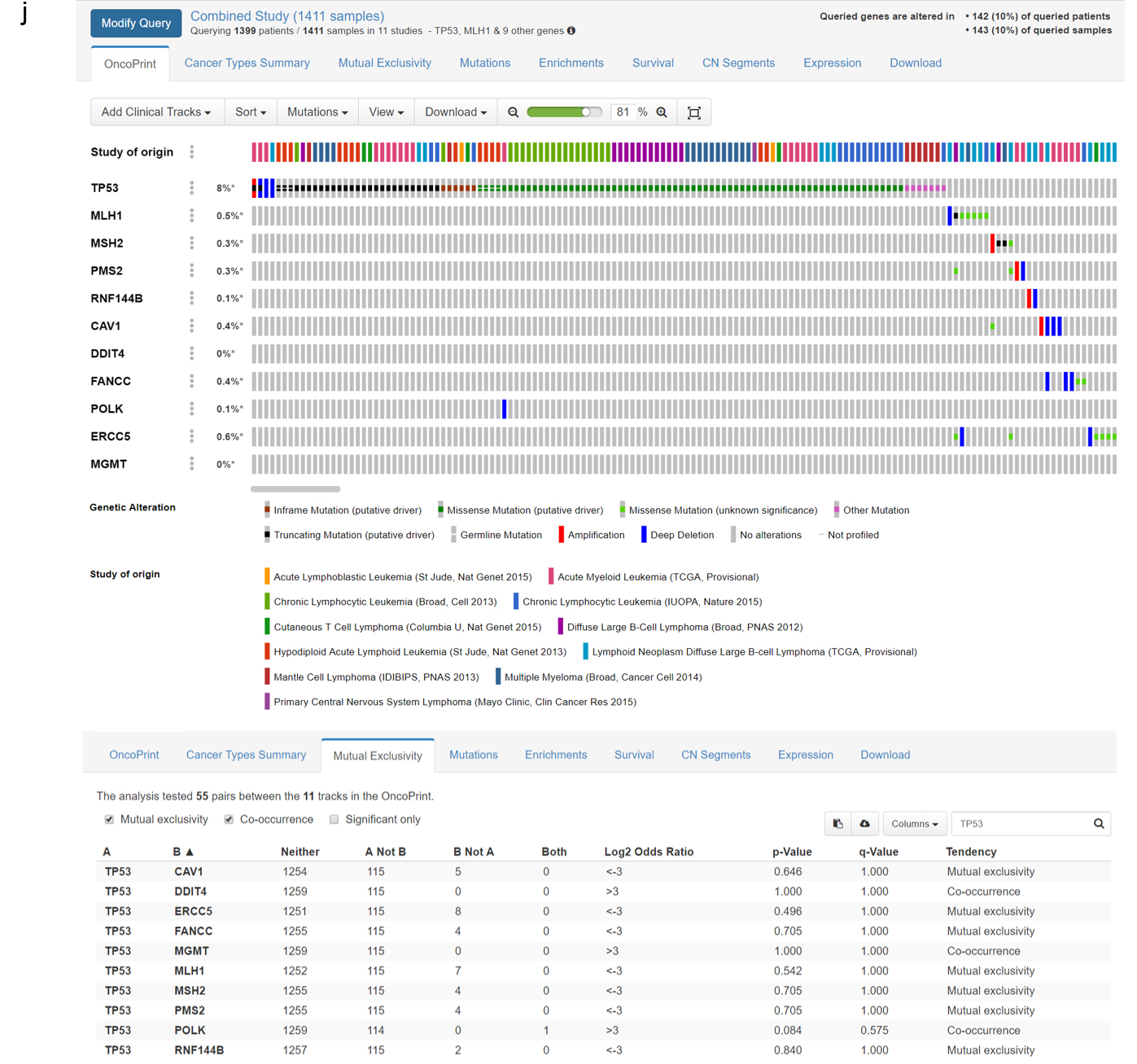

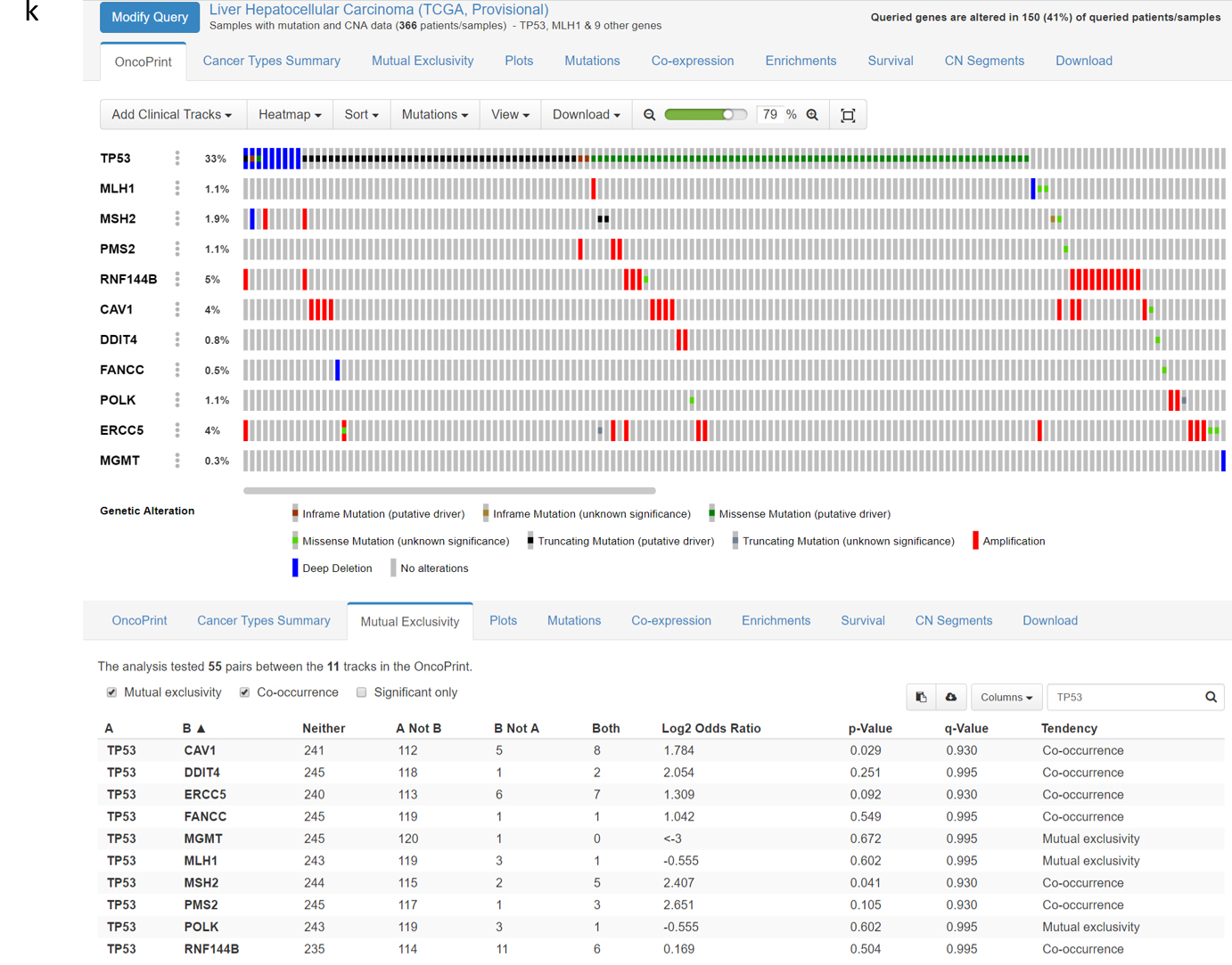

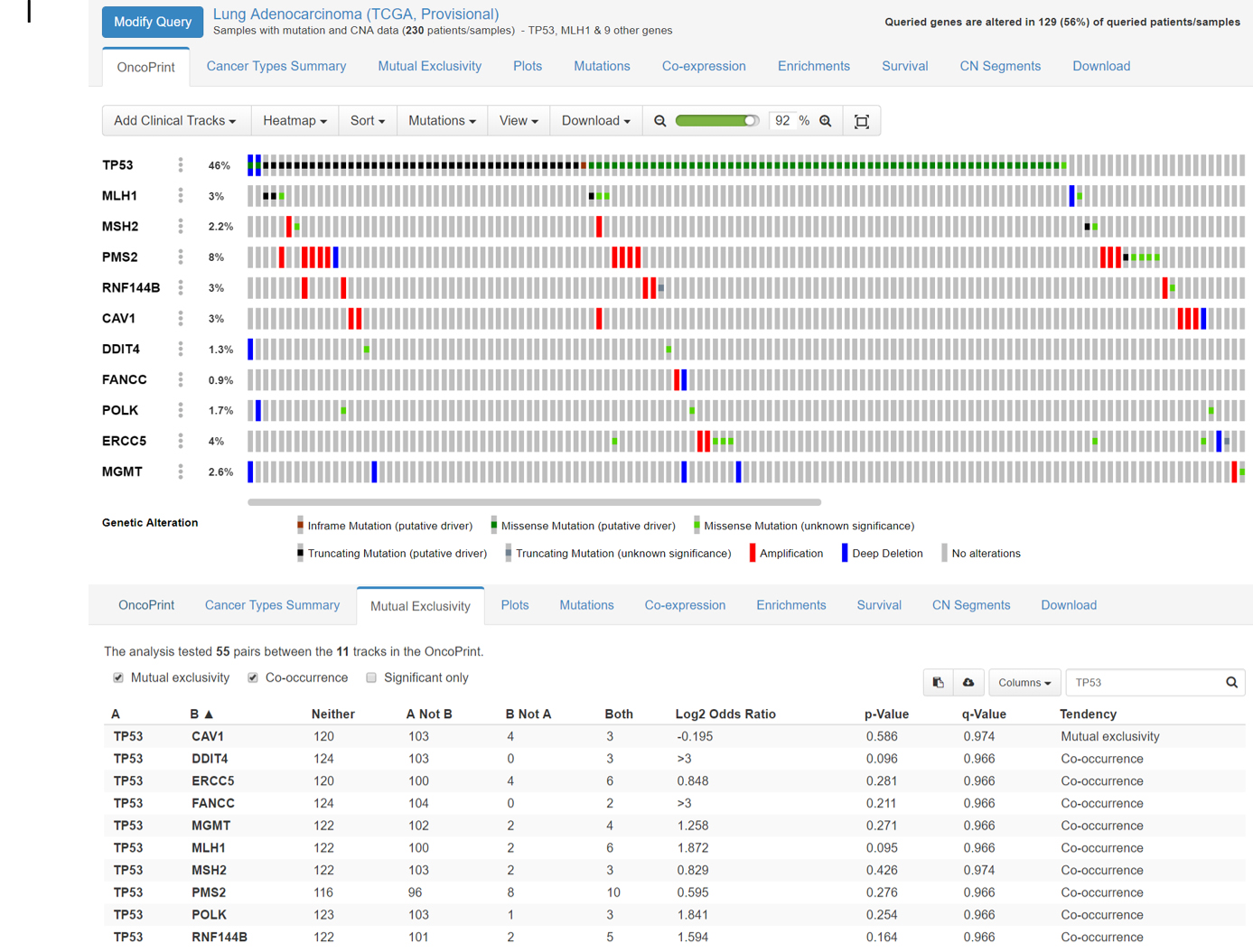

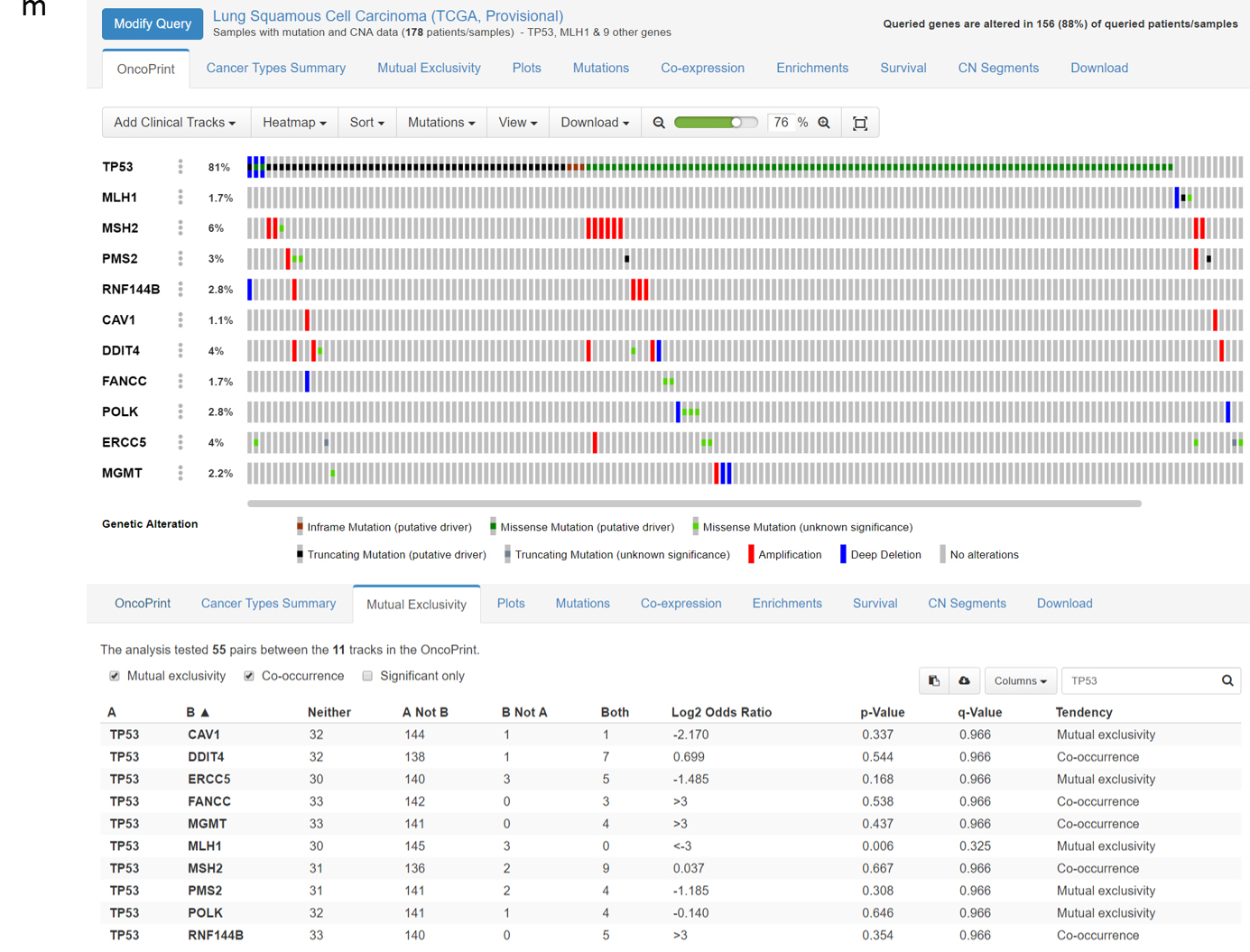

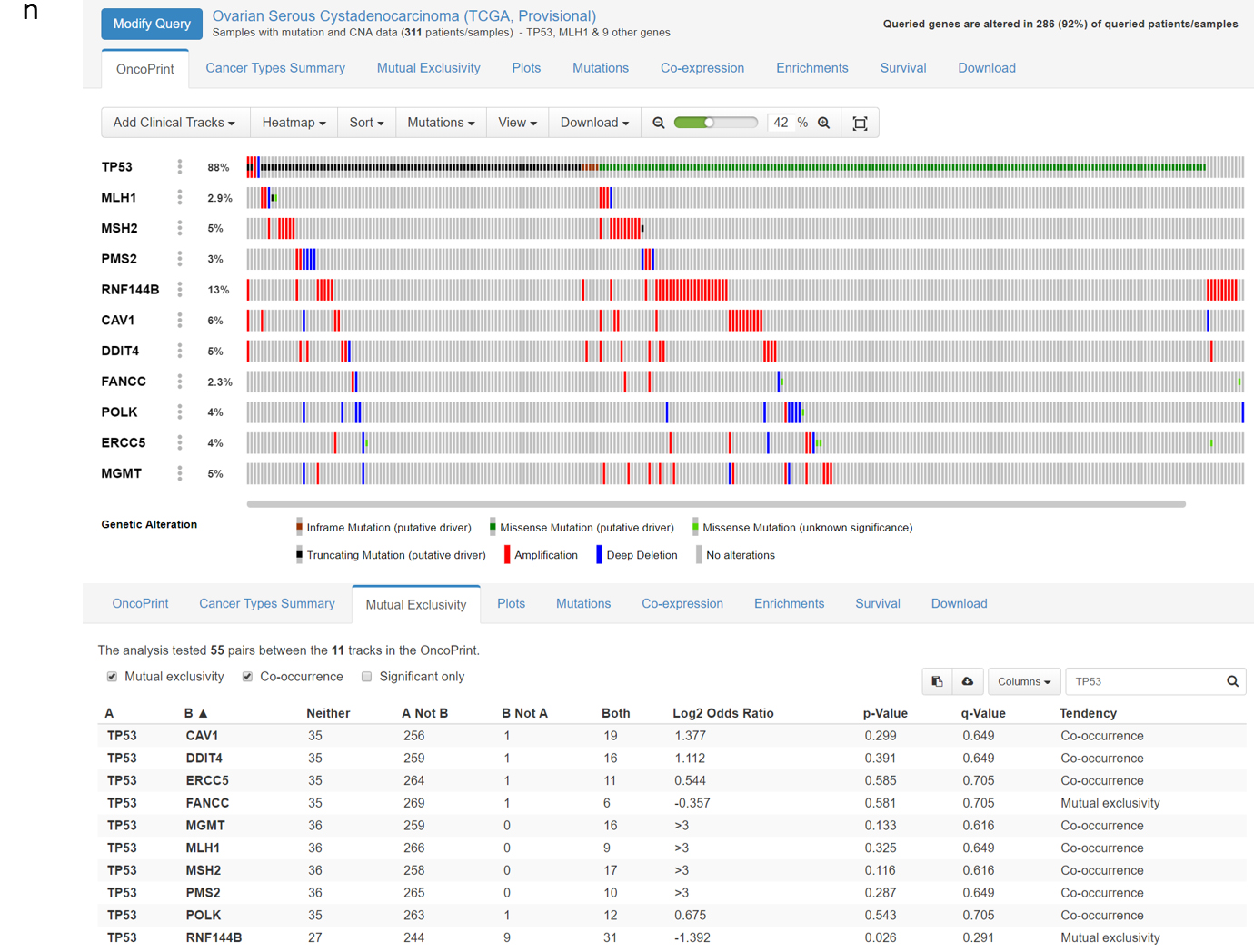

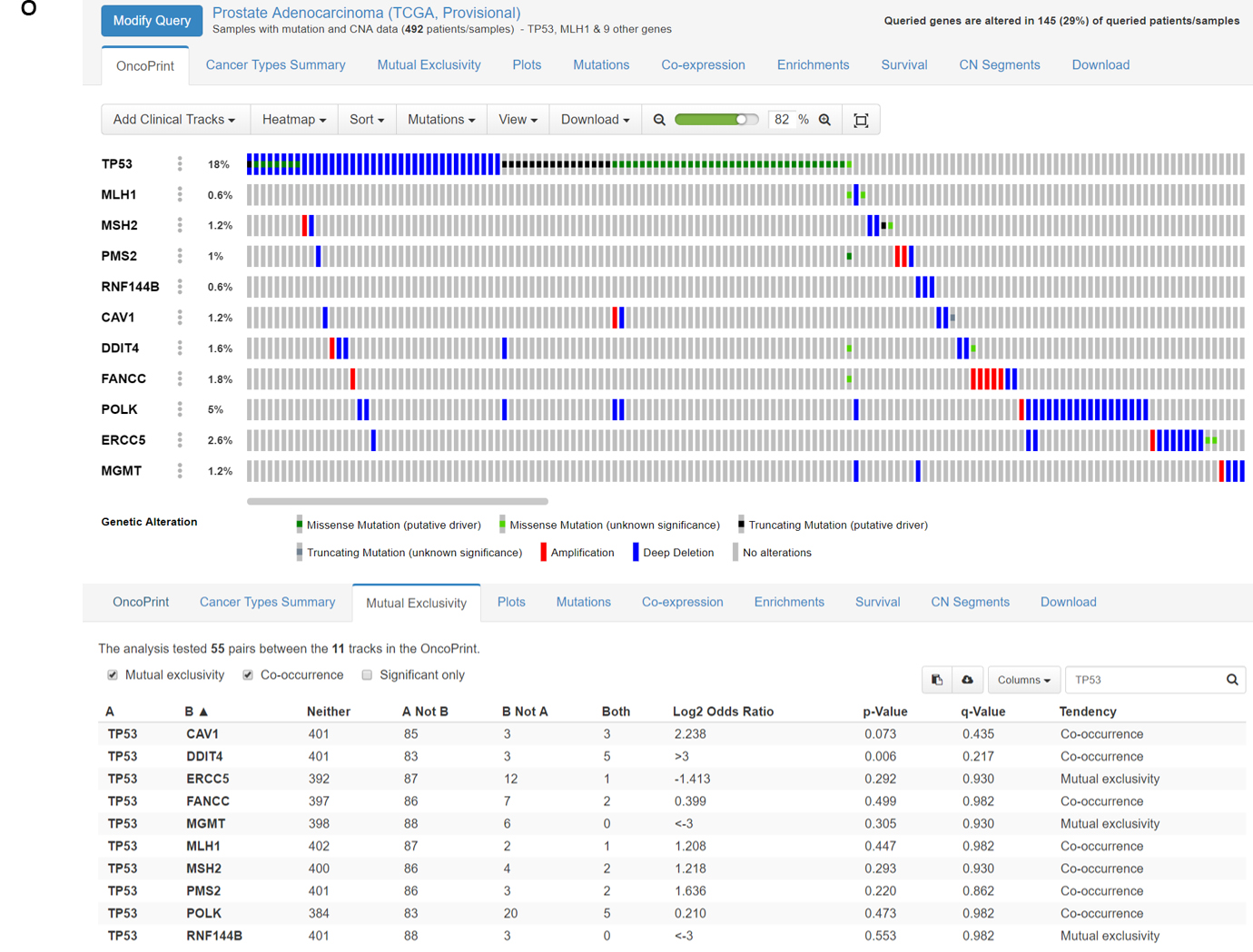

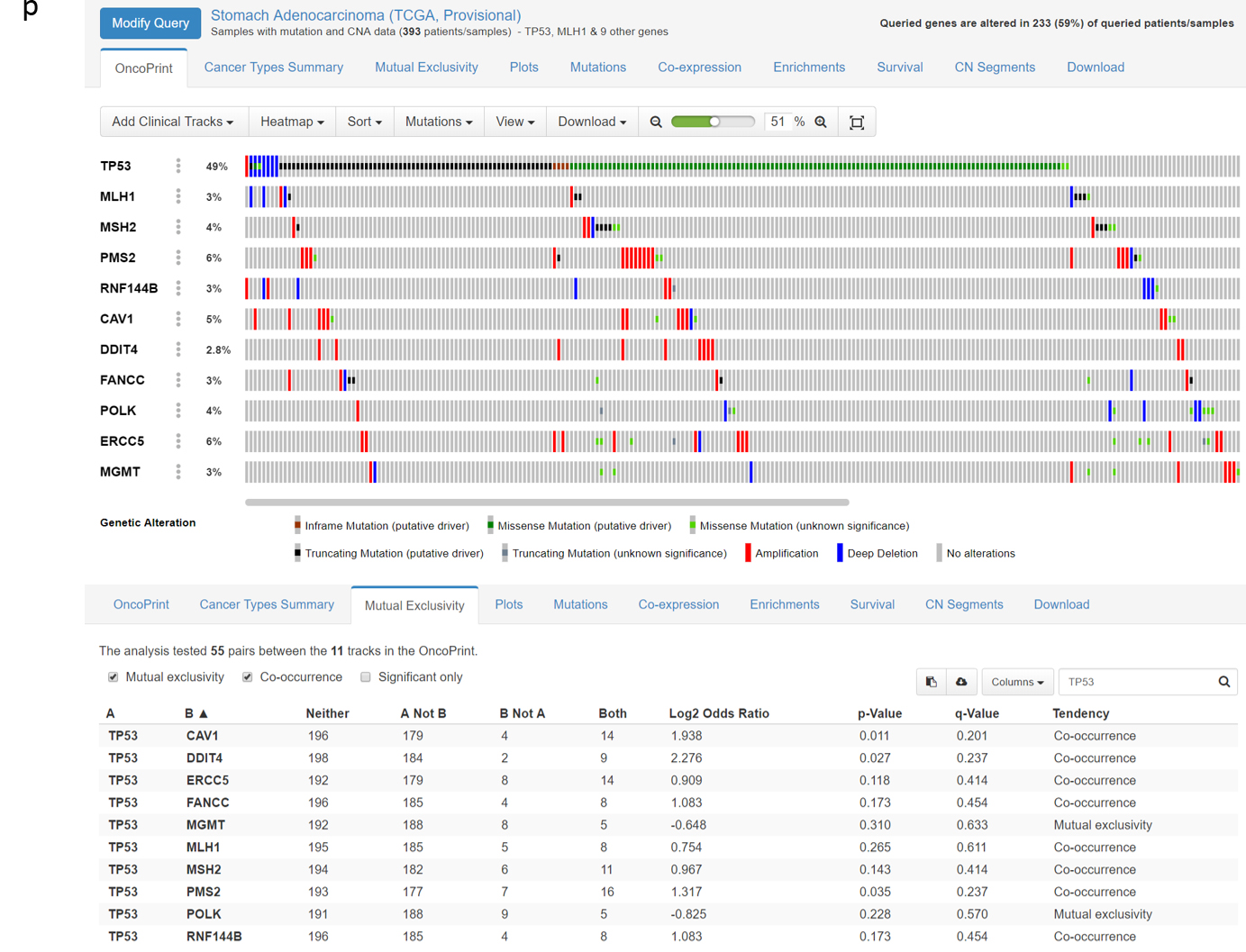

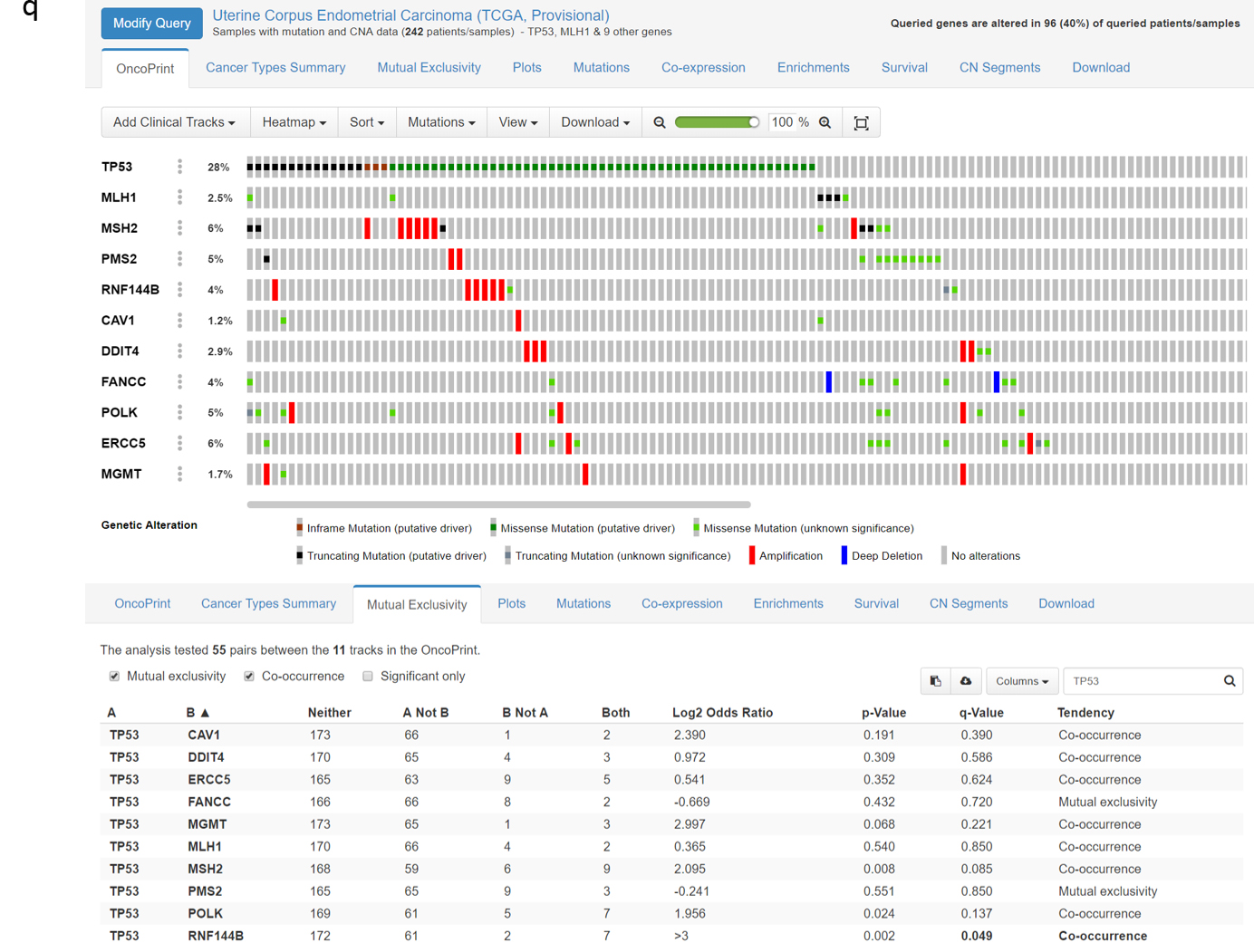

**Supplementary Figure 1. *P53(TP53)* and DNA repair gene mutation distribution in human malignancies analyzed in the same way as in the publication by Janic A, et al.^8^ based on cBioPortal^11,12^ data. a.** colorectal cancer; **b.** melanoma; **c.** glioma; **d.** breast invasive carcinoma; **e.** Adrenocortical Carcinoma; **f.** cervical cancer; **g.** cholangiocarcinoma; **h.** esophageal adenocarcinoma; **i.** head and neck cancer; **j.** hematological malignancies; **k.** liver cancer; **l.** lung adenocarcinoma; **m.** lung squamous cell carcinoma; **n.** Ovarian cancer; **o.** prostate cancer; **p.** Stomach adenocarcinoma**; q.** endometrial carcinoma.

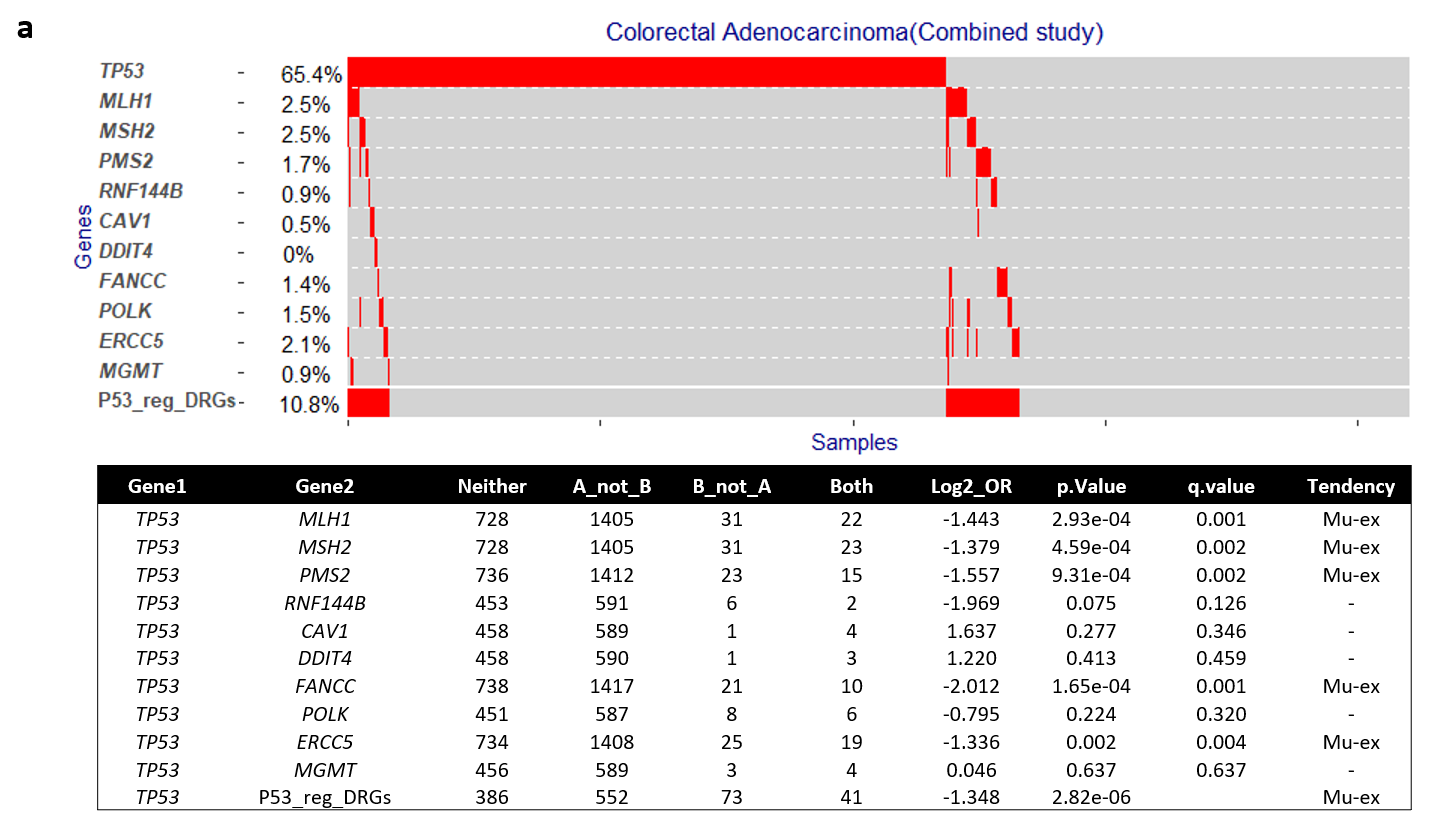

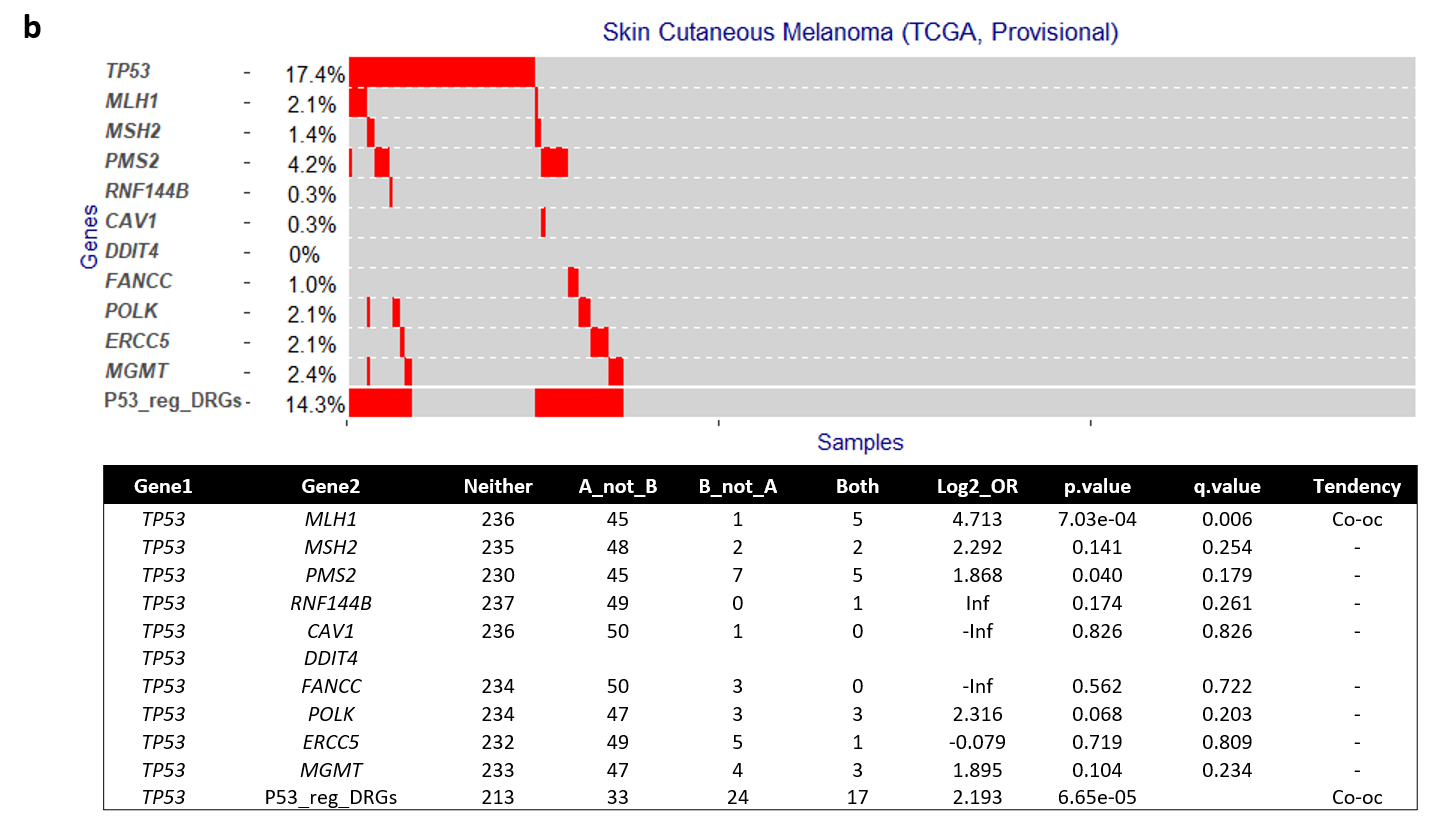

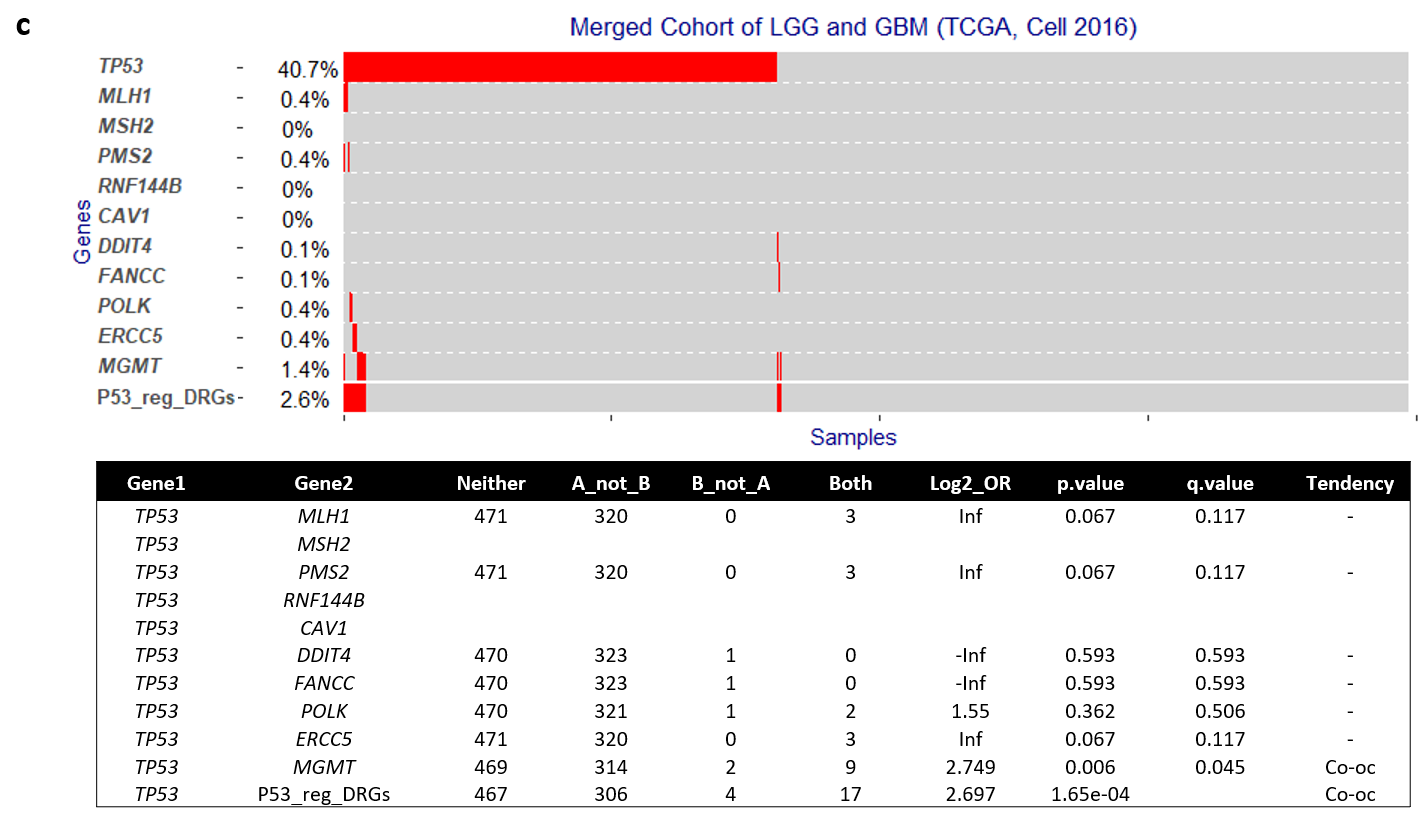

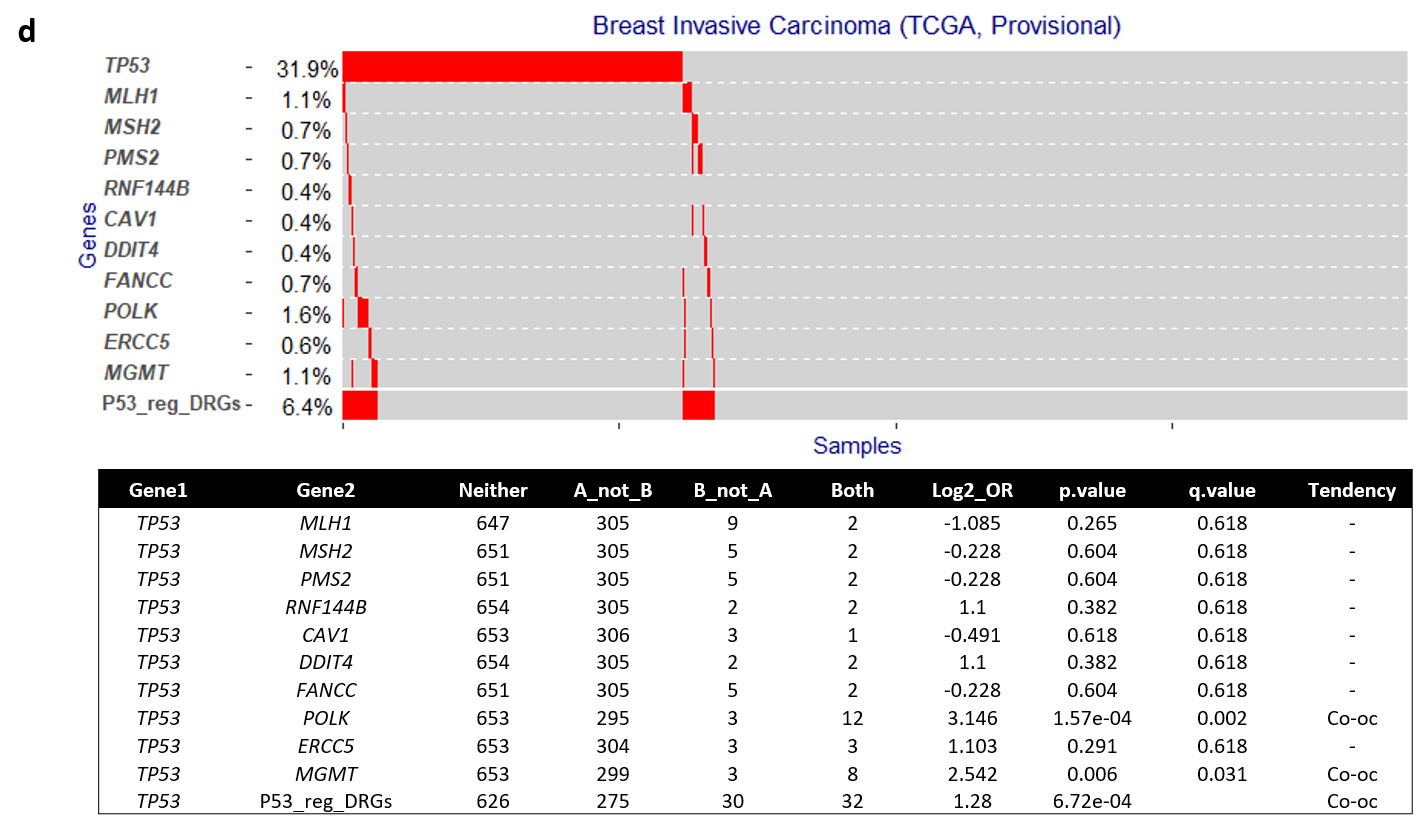

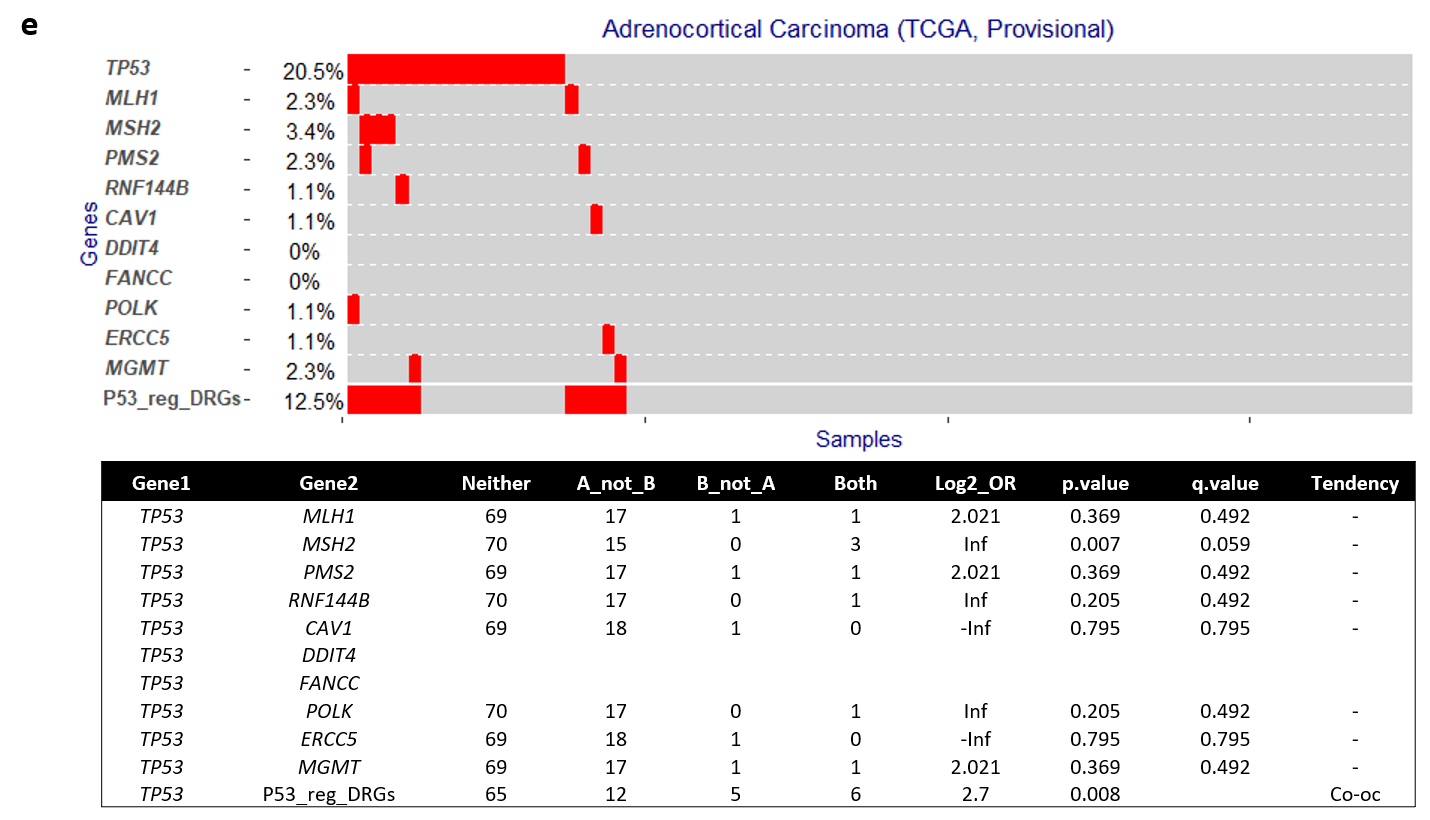

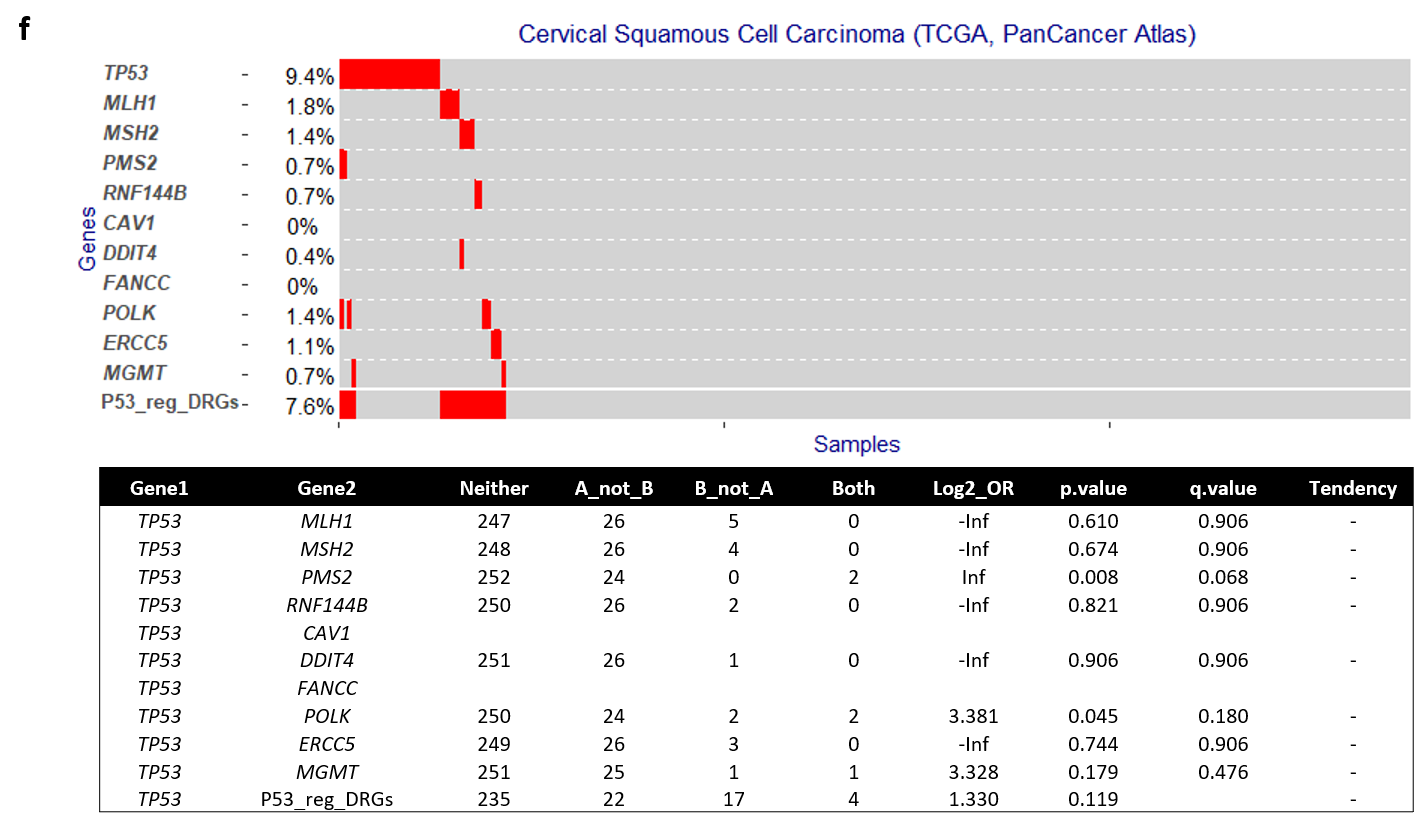

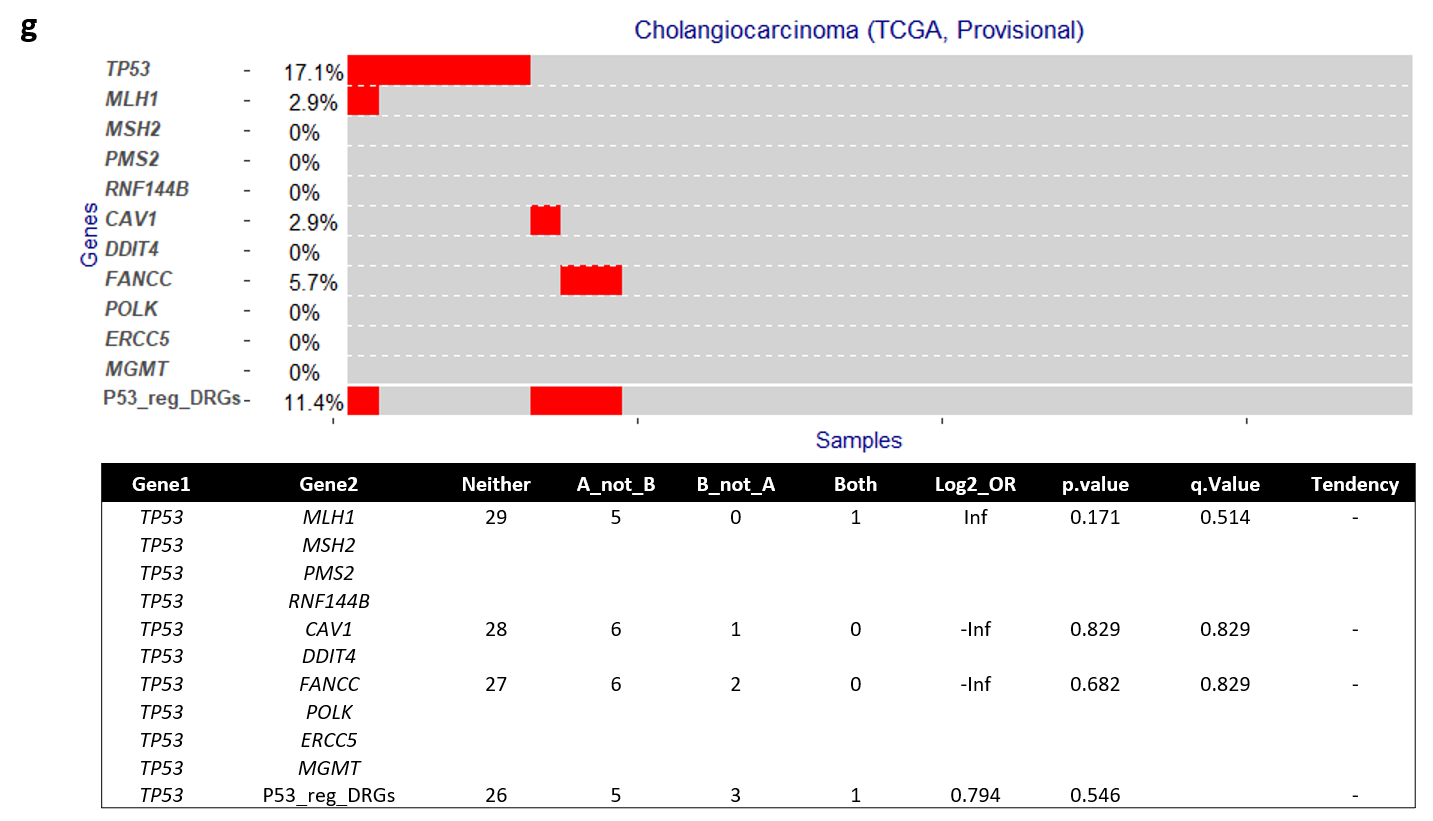

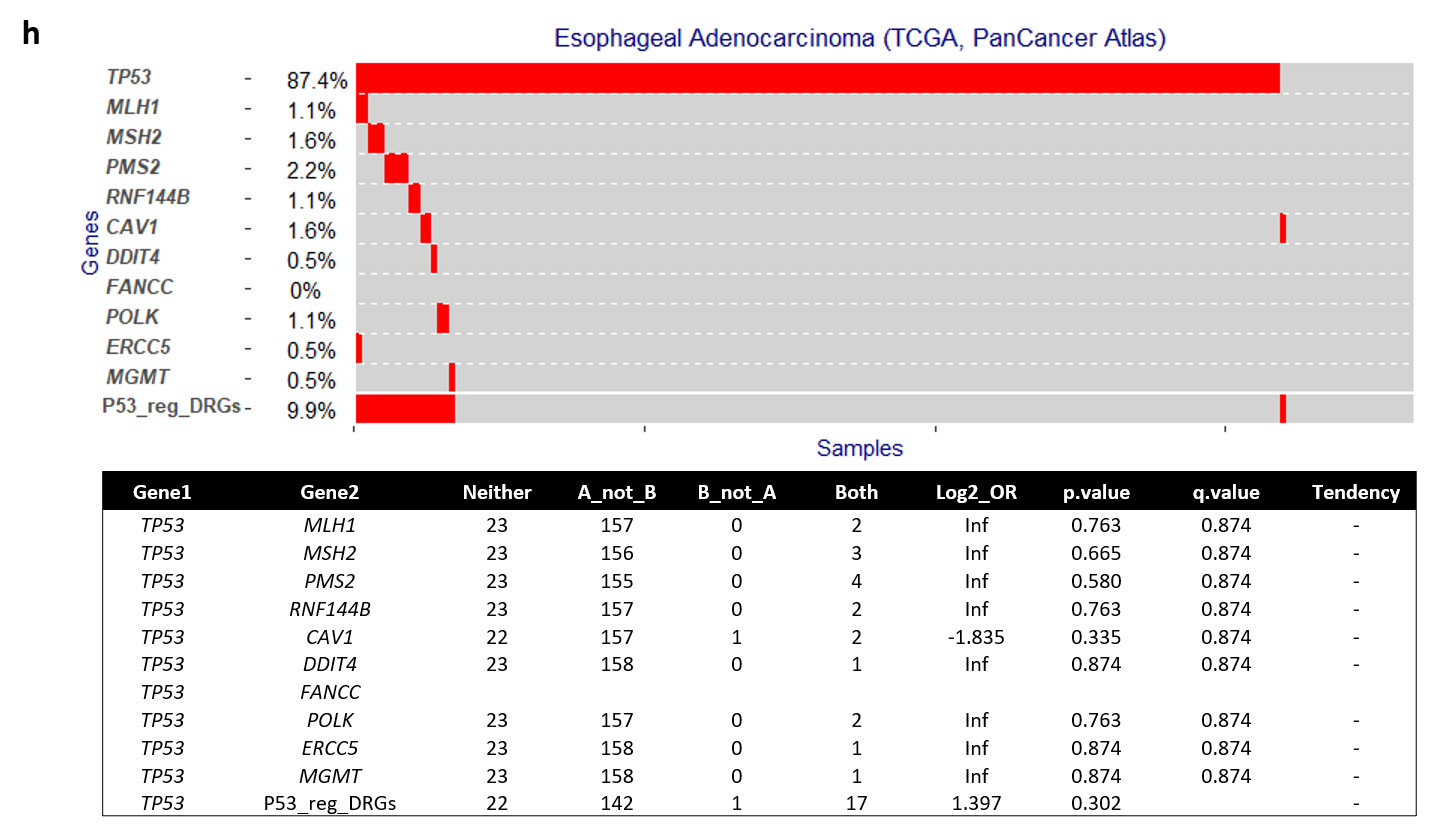

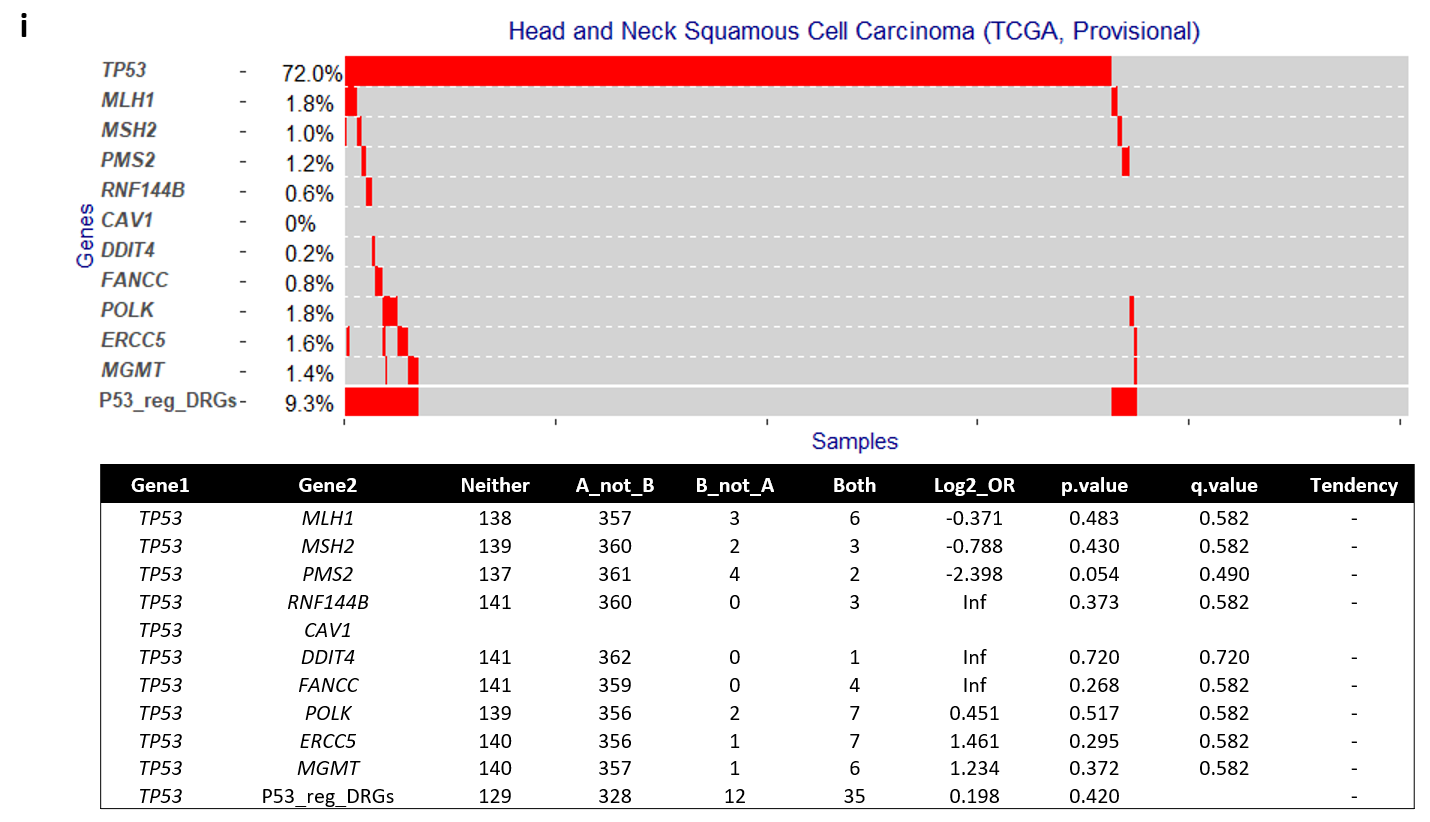

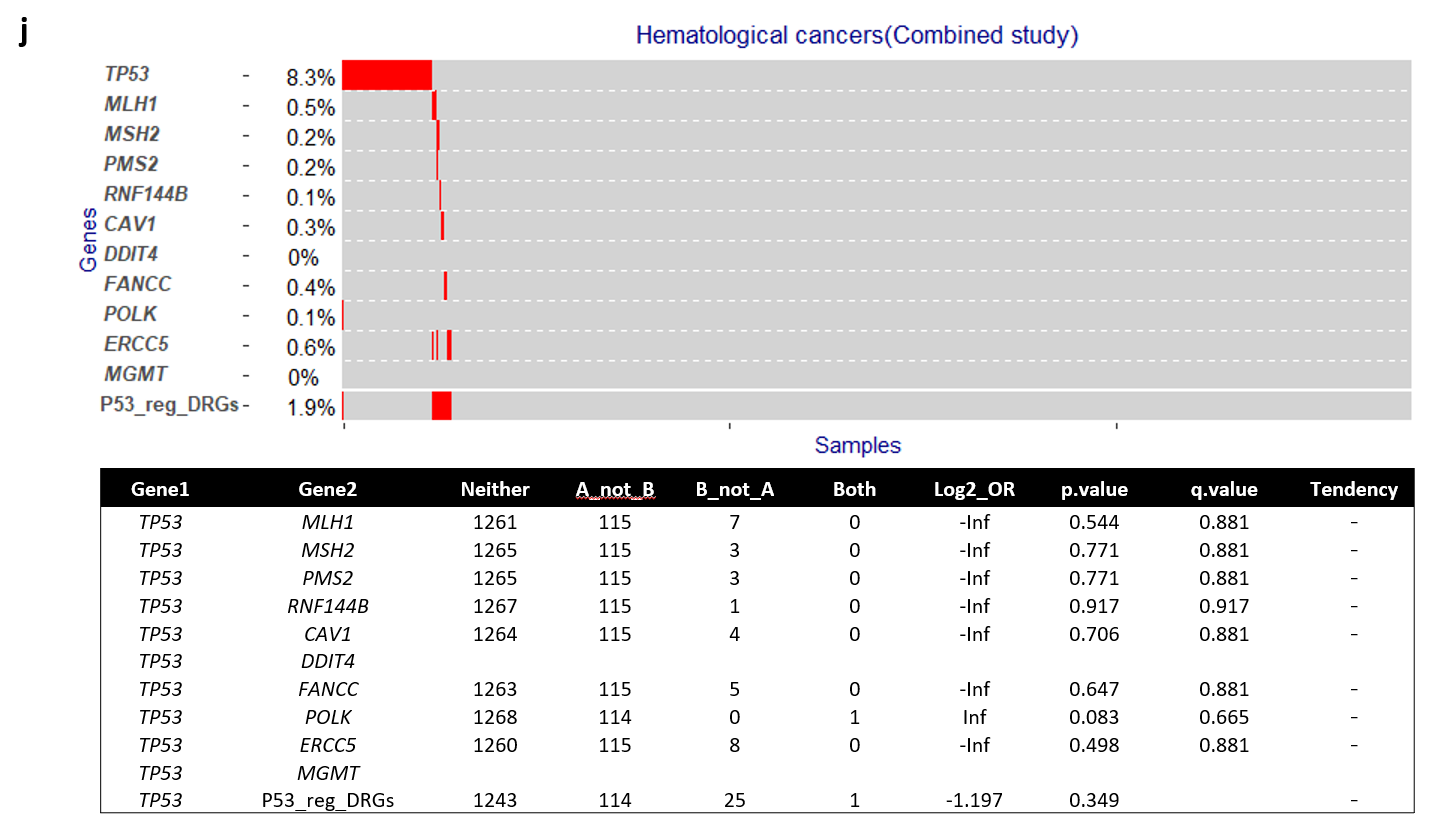

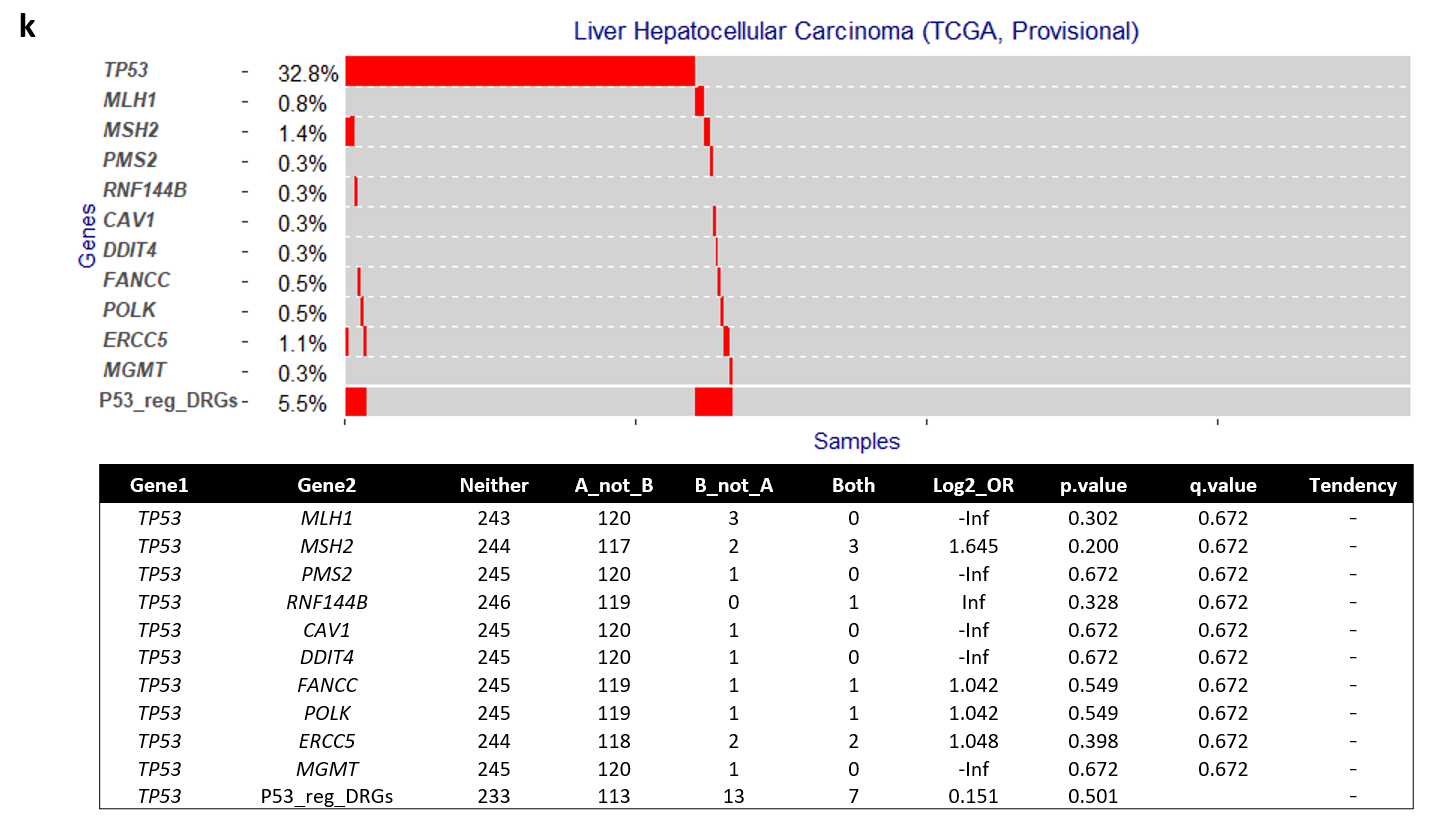

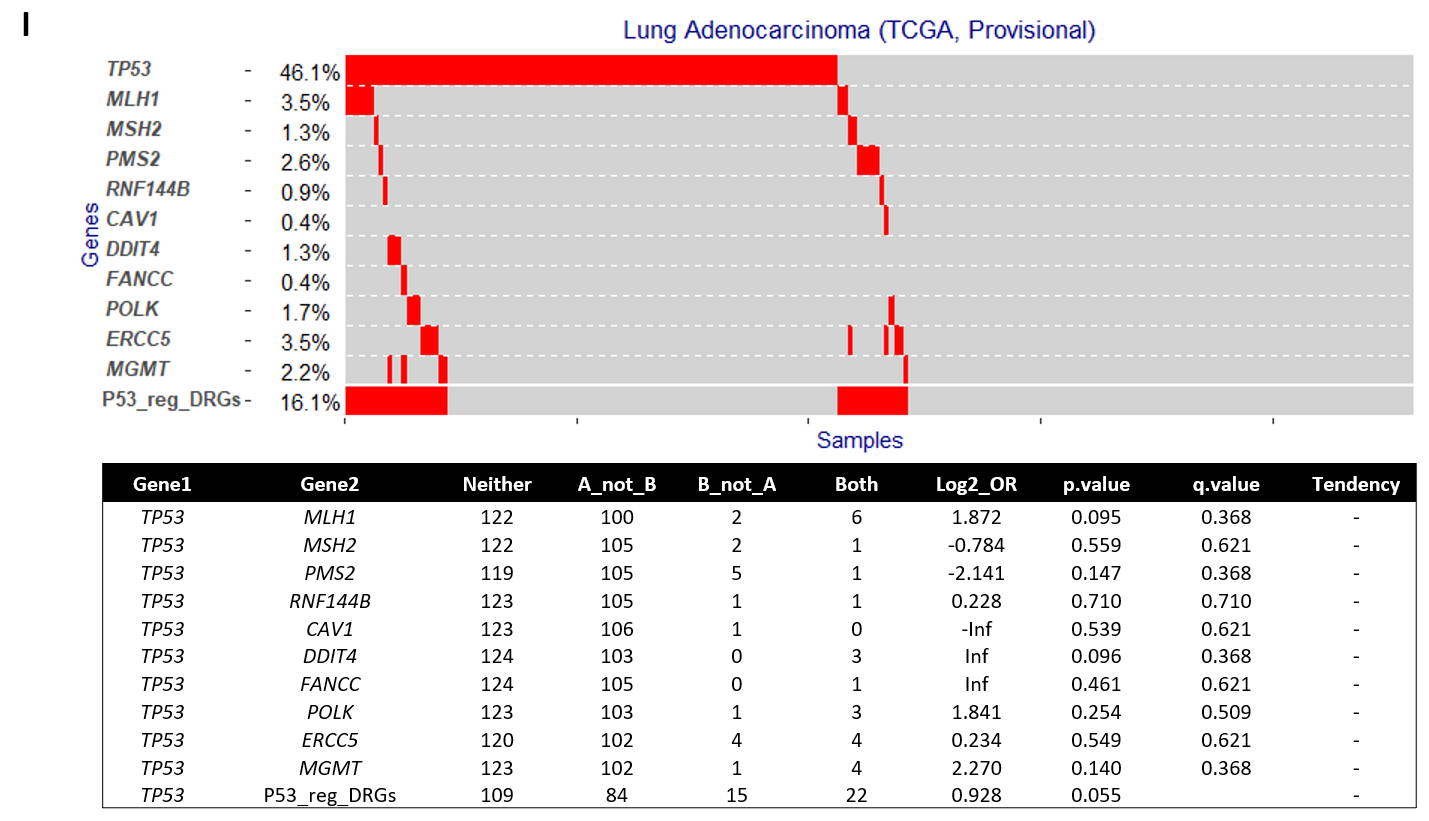

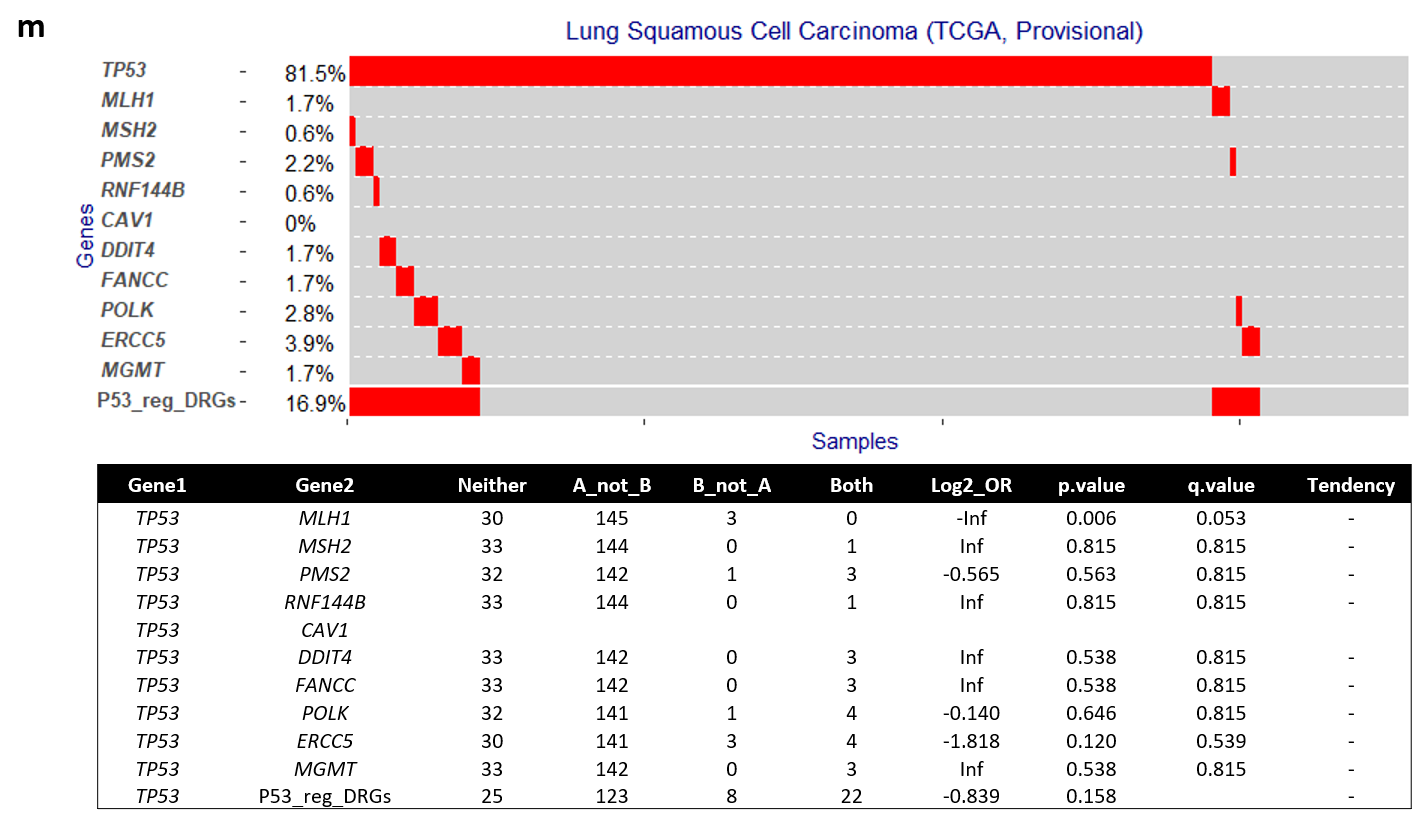

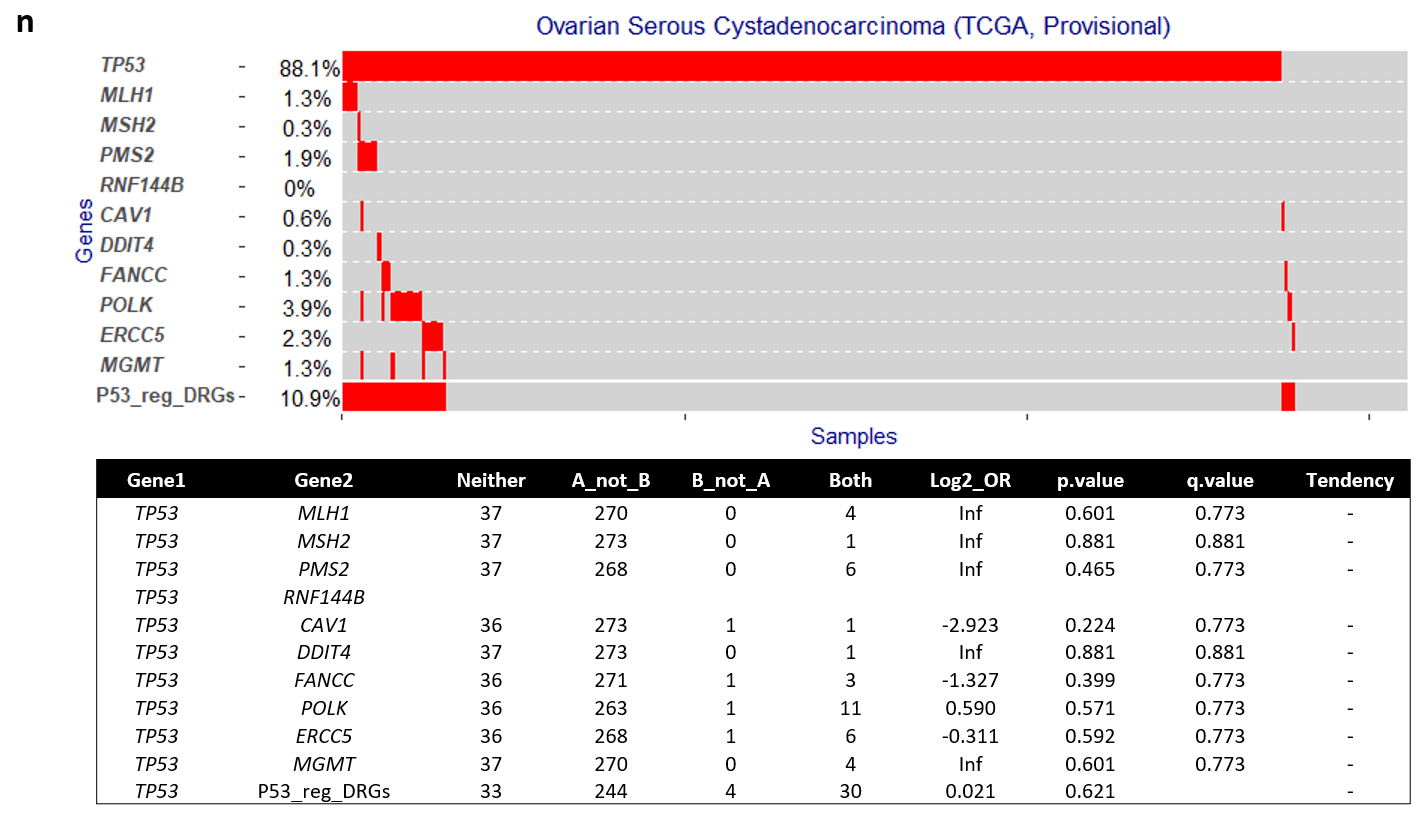

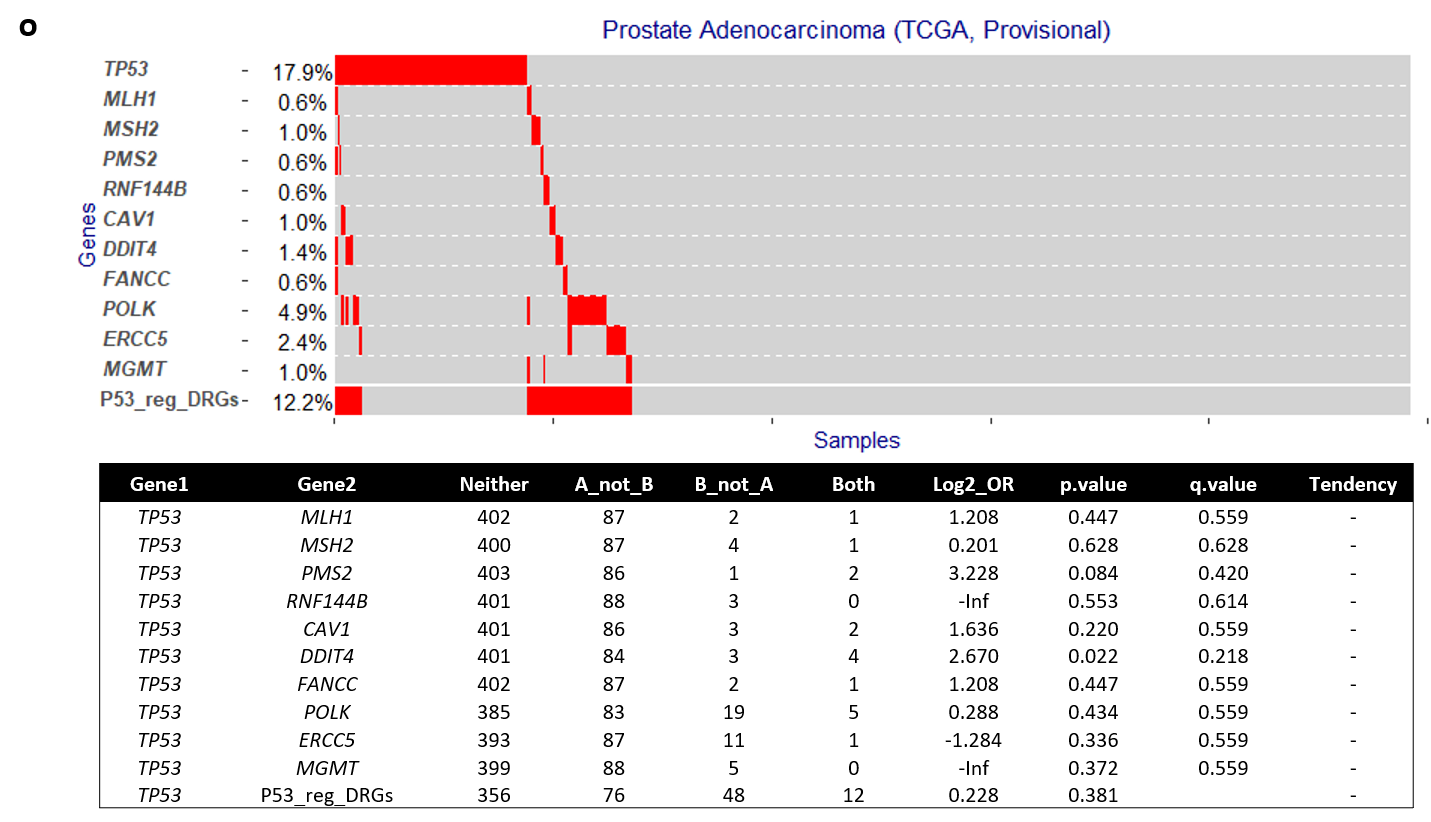

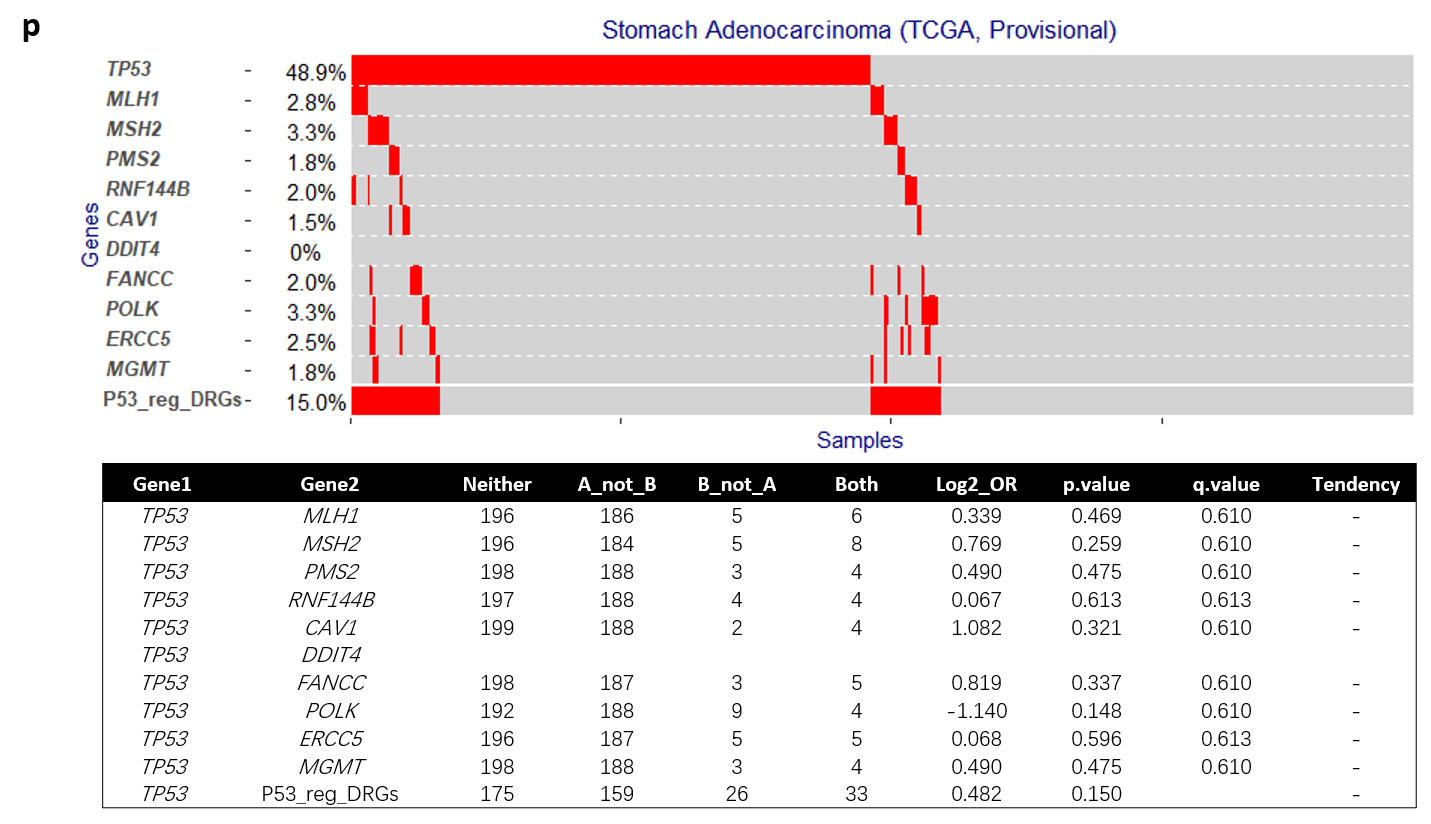

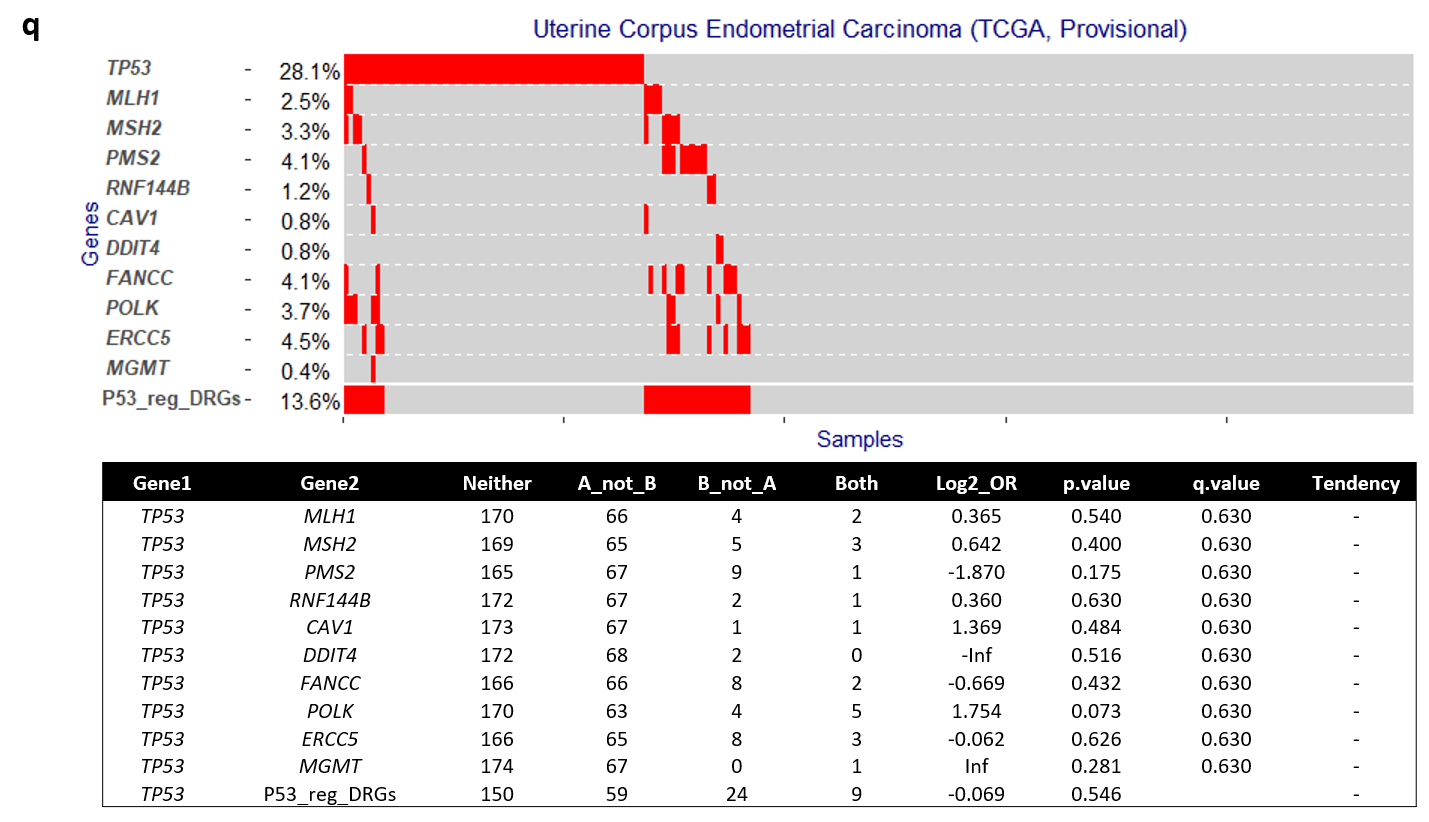

**Supplementary Figure 2. Distribution of *P53(TP53)* and DNA repair gene mutations (excluding amplification) in human malignancies based on cBioPortal^11,12^ data. a.** colorectal cancer; **b.** melanoma; **c.** glioma; **d.** breast invasive carcinoma; **e.** Adrenocortical Carcinoma; **f.** cervical cancer; **g.** cholangiocarcinoma; **h.** esophageal adenocarcinoma; **i.** head and neck cancer; **j.** hematological malignancies; **k.** liver cancer; **l.** lung adenocarcinoma; **m.** lung squamous cell carcinoma; **n.** Ovarian cancer; **o.** prostate cancer; **p.** Stomach adenocarcinoma**; q.** endometrial carcinoma. P53_reg_DRGs: mutation in any of the 10 p53 regulated DNA repair genes in combination; e-0n: X10^-n^; OR: odd ratio; Mu-ex: mutual exclusivity; Co-oc: co-occurrence; Inf: Infinity.
